# Transferability of ion force fields to OPC water: Maintaining single-ion and ion-pairing properties

**DOI:** 10.64898/2026.03.31.715553

**Authors:** Christian Wiebeler, Sebastian Falkner, Nadine Schwierz

**Affiliations:** Institute of Physics, University of Augsburg, 86159 Augsburg, Germany

**Author notes:** (Electronic mail).

## Abstract

Accurate ion force fields are essential for molecular dynamics simulations of biomolecular systems, particularly in combination with modern water models such as OPC. While OPC water provides an accurate description of key water properties and has been successfully applied in biomolecular simulations, the transferability of existing ion force fields to this model remains an open question. Here, we systematically assess the transferability of monovalent and divalent ion force field parameters (Li^+^, Na^+^, K^+^, Cs^+^, Mg^2+^, Ca^2+^, Sr^2+^, Ba^2+^, Cl^−^ and Br^−^) to OPC water by comparing single-ion and ion-pairing properties with experimental data. Our analysis reveals that no single literature parameter set provides accurate results for all ions when directly transferred to OPC water. We hence introduce the MS/G-LB(OPC) force field, which combines Mamatkulov-Schwierz-Grotz cation parameters with Loche-Bonthuis anion parameters. MS/G-LB(OPC) reproduces hydration free energies, first-shell structural properties and activity derivatives at low salt concentrations. Our results demonstrate that transferring ion parameters to OPC can lead to significant and ion-specific deviations from experimental data, making careful validation essential. At the same time, the systematic transfer and combination of ion parameters from existing force fields can provide a practical and computationally efficient alternative to full reparameterization. MS/G-LB(OPC) is available at https://git.rz.uni-augsburg.de/cbio-gitpub/opcion-force-fields.

## I. INTRODUCTION

Over the past decade, the four-site OPC water model has been increasingly adopted in biomolecular simulations due to its improved description of bulk water properties^1^. Since water constitutes the major component of typical biomolecular simulation systems, both the accuracy and the computational efficiency of the employed water model are of central importance. In combination with modern biomolecular force fields, OPC has been shown to yield improved agreement with experimental observables for both proteins and nucleic acids^2,3^, including more realistic conformational ensembles of intrinsically disordered proteins^4,5^ and enhanced agreement with NMR and structural data for RNA systems^6,7^. In addition, OPC reproduces liquid-water electrostatics with high fidelity^1^ and improves simulations of DNA^8,9^ and ligand binding free energies^10^, motivating its increasing use in molecular dynamics (MD) simulations of protein and nucleic acids^11^.

Ions are an essential component of biomolecular simulations, as they control electrostatic screening, stability, and specific interactions with proteins and nucleic acids^12–15^. With the growing adoption of OPC water, an accurate description of ionic solvation and ion-ion interactions in this solvent has therefore become increasingly important. It is tempting to combine OPC water with established ion force fields in order to leverage the strengths of each parameter set. However, even for simple monovalent and divalent ions, transferring parameters between water models can lead to substantial changes in solvation structure, thermodynamic properties, and ion-ion interactions^16–18^. It is therefore crucial to assess whether the transfer of ion parameters to OPC water yields physically meaningful and transferable results.

This raises the question of whether ion force fields must be optimized from scratch for each water model or whether parameter transfer is still possible. Döpke *et al*.^17^ showed that ion parameters optimized for TIP4P/Ew remain transferable to TIP4P/2005, yielding accurate thermodynamic properties without reparameterization. Loche et al.^19^ developed a global optimization procedure for Na^+^, K^+^, Cl^−^ and Br^−^ with transferability to TIP3P, TIP4P/*ε*, and, to a lesser degree of accuracy, TIP4P. This work was recently extended to Li^+^ and Cs^+^.^20^ In previous work^16,21^, we demonstrated that Mg^2+^ force field parameters originally optimized in combination with SPC/E water reproduce a broad range of experimental ion properties when transferred to OPC. The success of these approaches motivates the question whether similar transferability can be achieved for a broader set of ions in OPC water.

Monovalent cations such as Li^+^, Na^+^, K^+^, and Cs^+^ are ubiquitous in electrolyte solutions, cellular environments, and ion-specific biochemical processes^13,22^, while divalent cations such as Ca^2+^, Sr^2+^, and Ba^2+^ play essential roles in signaling, structural stabilization, and catalysis^23,24^. Likewise, anions such as Cl^−^ and Br^−^ play a key role in hydration, ion pairing, and electrostatic screening in electrolyte solutions^25^. Accurate force field parameters for these ions are therefore indispensable for biomolecular simulations^26^.

In this work, we extend the transferability analysis^16^ previously introduced for Mg^2+^ to a comprehensive set of monovalent (Li^+^, Na^+^, K^+^, Cs^+^) and divalent (Mg^2+^, Ca^2+^, Sr^2+^, Ba^2+^) cations together with the anions Cl^−^ and Br^−^. We examine the transferability of three ion force fields from the literature^19,27^ and assess their performance in OPC water with respect to key single-ion properties, including the first-shell hydration number, the first-shell ion-water distance, and the hydration free energy, as well as ion-pairing properties quantified by activity derivatives.

In contrast to our previous findings for Mg^2+^, where SPC/E-optimized ion parameters exhibited transferability to OPC^16^, the present systematic assessment reveals that no single cation-anion parameter set is sufficient to accurately describe all ions considered here. Instead, we identify a mixed anion/cation parameter combination consisting of Mamatkulov-Schwierz-Grotz cation parameters^29^ and Loche-Bonthuis anion parameters^19^, denoted MS/G-LB(OPC), which provides the most accurate reproduction of experimental single-ion and ion-pairing properties. Notably, MS/G-LB(OPC) reproduces experimental observables with an accuracy comparable to, and for single-ion properties exceeding, that of the OPC-optimized ion force fields^30,31^.

Our results demonstrate that the transfer and combination of ion force field parameters optimized against experimental single-ion and ion-pairing data provide a viable alternative to full reparameterization. At the same time, careful validation remains essential, as force field parameter transfer can lead to non-trivial and ion-specific effects.

## II. METHODS

### Force fields

The pair potential between atoms *i* and *j* is described as the sum of the Coulomb and the 12-6 Lennard-Jones (LJ) potential,

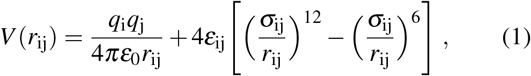

where *q*_i_ is the charge, *r*_ij_ the distance, and *ε*_0_ the dielectric constant of vacuum. The LJ term contains the interaction strength *ε*_ij_ and the diameter *σ*_ij_. Interactions between different atom types are described using the unmodified Lorentz-Berthelot combination rules:

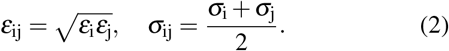

For the *ε*_i_ and *σ*_j_ the following parameters were used: Parameter set 1 (Mamatkulov–Schwierz^29^) was optimized with the TIP3P^32^ water model. Parameter set 2 (Fyta–Netz^27,28^) was optimized with the SPC/E^33^ water model. Both sets were designed to reproduce experimental solvation free energies, ion–water distances, and activity derivatives. Parameter set 3 (Loche–Bonthuis for Na^+^, K^+^, Cl^−^ and Br^−^ and Herrera-Scalfi for Li^+^, Cs^+19,20^ was obtained by a global optimization procedure simultaneously optimizing the solvation free energy and activity derivatives. In addition, we compare the Li–Merz^30,31^ hydration free energy (HFE) parameter set, which was optimized with the OPC water model (set 4).

In our previous work, we found that the original MS force field significantly underestimated the water exchange rate in the first hydration shell of Mg^2+^ compared to experimental measurements.^34^ To address this limitation, the microMg force fields were developed and parameterized to reproduce experimental water exchange kinetics while simultaneously maintaining accurate hydration free energies, hydration-shell structure, and activity derivatives for different water models including OPC^16,21,35^. Therefore, in the final MS/G-LB(OPC) force field (Table I), we employ the microMg(OPC) parameters for Mg^2+^ in OPC water^16^.

**TABLE I.** MS/G-LB(OPC) force field parameters. *σ*_*ii*_ and *ε*_*ii*_ denote the ion-ion Lennard-Jones parameters, while *σ*_*io*_ and *ε*_*io*_ denote the ion-water parameters. The parameters are to be used with Lorentz-Berthelot combination rules.

| | $\sigma_{ii}$ (nm) | $\epsilon_{ii}$ (kJ/mol) | $\sigma_{io}$ (nm) | $\epsilon_{io}$ (kJ/mol) |
| --- | --- | --- | --- | --- |
| Li<sup>+</sup> | 0.151813 | 0.604036 | 0.234234 | 0.733355 |
| Na<sup>+</sup> | 0.260719 | 0.114553 | 0.288687 | 0.319364 |
| K<sup>+</sup> | 0.298839 | 0.604036 | 0.307747 | 0.733355 |
| Cs<sup>+</sup> | 0.354915 | 0.604036 | 0.335785 | 0.733355 |
| Mg<sup>2+</sup> | 0.103643 | 290.584188 | 0.210149 | 16.084916 |
| Ca<sup>2+</sup> | 0.226453 | 2.338830 | 0.271554 | 1.443053 |
| Sr<sup>2+</sup> | 0.282385 | 1.056590 | 0.299520 | 0.969920 |
| Ba<sup>2+</sup> | 0.354615 | 0.277190 | 0.335635 | 0.496789 |
| Cl<sup>-</sup> | 0.430000 | 0.420000 | 0.373328 | 0.611515 |
| Br<sup>-</sup> | 0.443000 | 0.750000 | 0.379828 | 0.817172 |

### Solvation free energy, radius, and coordination number

We calculated the solvation free energy of neutral ion pairs using free energy perturbation (FEP) simulations. Finite size corrections^36^ and a pressure correction term^37^ were applied. Note that the interfacial crossing^37^ cancels for neutral ion pairs. In addition, we calculated the ion-water oxygen distance *R*_1_ and the coordination number *n*_1_ as structural descriptors of the first hydration shell.

All simulations were performed at a temperature of 300 K, consistent with previous ion force field optimization studies. Experimental reference data reported for 298.15 K were therefore corrected to 300 K following the procedure introduced in Ref. 19. An overview of the simulation setups employed, more details on the determination of the solvation free energy and further results are reported in the Supporting Information (SI).

### Activity derivative

We calculated the activity derivatives for four different salt concentrations: 0.26 m, 0.52 m, 1.06 m, and 2.12 m. The activity derivatives were obtained from NPT simulations and Kirkwood-Buff theory^38,39^ (see SI for more details). Since the experimental activity derivatives are reported as a function of molality rather than molarity, the simulation results are also reported as function of molality.

### Self-diffusion coefficient and ion-water potential of mean force

We calculated the self-diffusion coefficient *D* and the ion-water potential of mean force (PMF). *D* was calculated from the slope of the mean-square displacement obtained in 50 ns NVT simulations of single ions in OPC water. The diffusion coefficient was corrected for system size effects^40^ and rescaled with the ratio of water model and experimental viscosity^41^. This procedure compensates for systematic deviations in the solvent dynamics and is typically used to isolate the contribution of the Mg^2+^ force field. For the OPC water model, however, this correction is relatively minor because its viscosity at 298 K (*η≈*0.80 mPa *·*s)^42^ is already close to the experimental value of water (*η* = 0.891 mPa*·* s).^41^ Finally, we calculated the ion-water oxygen radial distribution functions *g*(*r*) and obtained the PMF from Boltzmann inversion using 1 µs simulations in 1 m salt solutions. For Mg^2+^, we used umbrella sampling as in our previous work^16^.

### MD simulation setup

All MD simulations were performed using GROMACS version 2020.7^43^. We employed a cutoff of 1.2 nm for Coulomb and Lennard-Jones interactions. Long-range dispersion corrections for energy and pressure were applied to account for interactions beyond the cutoff. We used the particle-mesh Ewald method^44^ and a Fourier spacing of 0.12 nm with grid interpolation up to order 4. Typically, a timestep of 1 fs was utilized during equilibration and 2 fs for production.

Energy minimization was performed by steepest descent with a maximum of 50,000 steps. This was followed by two equilibration runs of 0.5 ns in NVT and 1 ns in NPT. These simulations were realized at 300 K and 1 bar, using the Berendsen thermostat and barostat^45^. During the equilibration of systems with multiple ions, we used position restraints on all ions to ensure that the hydration shells can form properly and prevent the artificial formation of inner-sphere ion pairs.

In the production runs, no restraints were applied. During these simulations, temperature was controlled using the velocity-rescaling thermostat of Bussi et al.^46^ with a coupling time constant of *τ* = 0.1 ps and a reference temperature of 300 K. For production runs in the NPT ensemble, pressure was maintained at 1 bar using the Parrinello-Rahman barostat^47^ with a pressure coupling time constant of *τ*_p_ = 5 ps. Trajectory frames were written every 2 ps for subsequent analysis. An overview of the various setups used in our simulations and short descriptions on the calculation of the different structural and thermodynamic properties are given in the SI and in previous works^29,35^.

## III. RESULTS

In the following, we assess the performance of ion force fields from the literature when transferred to OPC water. By systematically comparing simulated ion properties to experimental reference data, we identify the parameter combination that provides the overall most accurate description of monovalent and divalent ions in OPC water (Table 1).

### A. Solvation free energy, ion-water distance, and coordination number

To assess the transferability of ion force fields to OPC water, we evaluate their performance with respect to three key single-ion properties: the solvation free energy, the radius of the first hydration shell, and the corresponding coordination number. Figure 1 shows the deviation of the solvation free energy ΔΔ*G*_solv_ from experiments for the different force fields. The TIP3P-optimized Mamatkulov-Schwierz parameters perform well in OPC for monovalent ions, but deviations are observed for divalent cations. The Loche–Bonthuis and Herrera-Scalfi parameters appear particularly promising, likely due to their global optimization strategy designed to produce water-model–independent force fields^19,20^. However, their applicability is currently limited to monovalent cations. Among the SPC/E-optimized Fyta-Netz parameters, good agreement is obtained for K^+^ and Cs^+^ in OPC, whereas the other cations show pronounced deviations. This is unexpected given the previously reported transferability of SPC/E-optimized Mg^2+^ parameters^16^. We conclude that parameter transfer between water models can produce ion-specific effects that are difficult to anticipate.

**FIG. 1.**
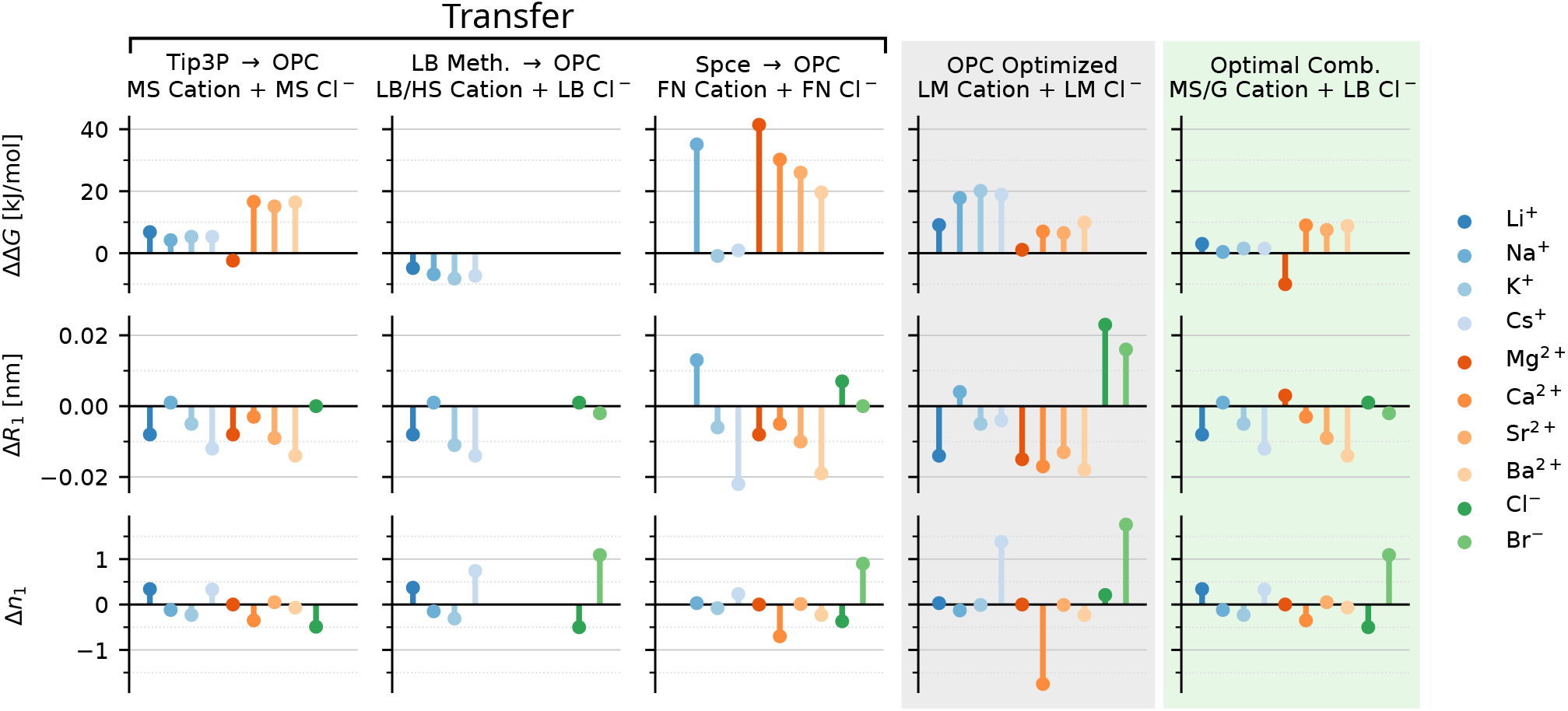
Difference between simulated and experimental single-ion properties: ΔΔ*G*_solv_ (top), Δ*R*_1_ (middle) and Δ*n*_1_ (bottom). Results for transferring three non-OPC-optimized parameter sets to OPC (white background). Results for the OPC-optimized Li-Merz parameters (grey background). Optimal combination of cation and anion parameter sets in MS/G-LB(OPC) (green background). The experimental values are from Refs. 48–50.

**FIG. 2.**
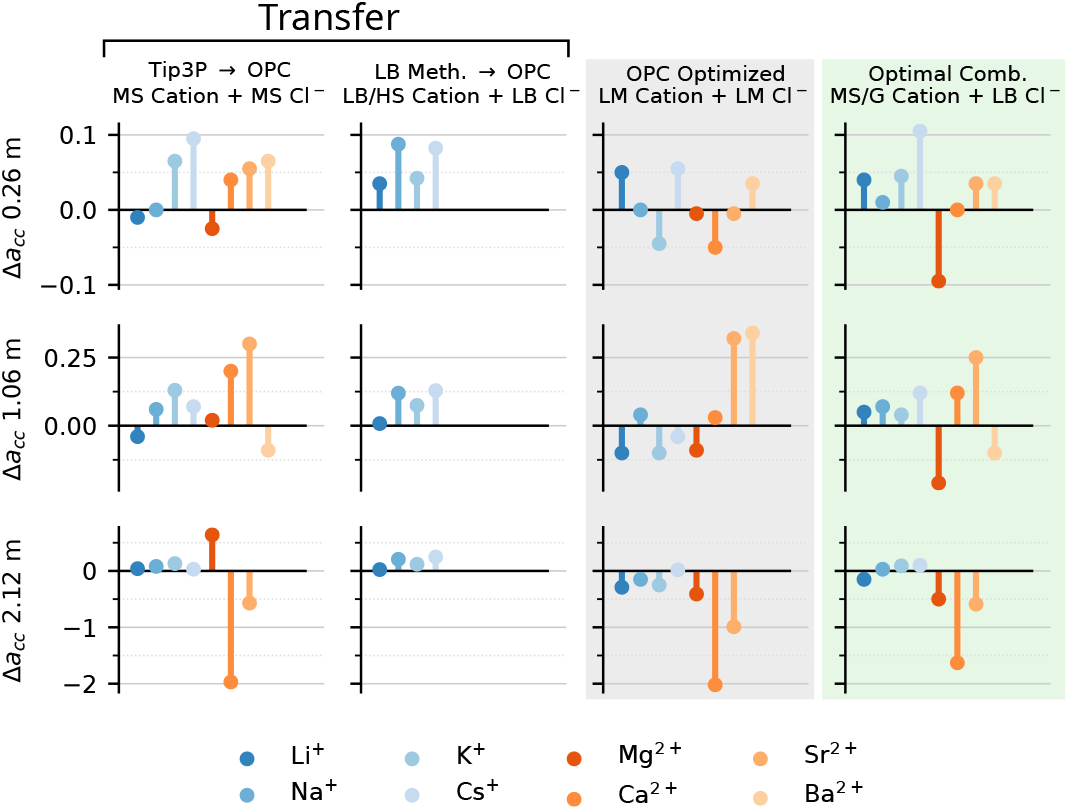
Difference between simulated and experimental activity derivatives: Δ*a*_*cc*_ at 0.26 m (top), at 1.06 m (middle) and 2.12 m (bottom). Experimental values were obtained from the data reported in Ref. [51].

Overall, there is no single literature parameter set that provides uniformly accurate results for all monovalent and divalent ions when directly transferred to OPC water. We hence performed a systematic evaluation of different cation-anion combinations. The best overall agreement with experimental data using the same reference anion and complete sets of 8 cations is obtained for the combination of Mamatkulov-Schwierz-Grotz cations with Loche-Bonthuis Cl^−^ (Figure 1, green background) and Br^−^ (see SI, Sections S8). We refer to this combined force field as MS/G–LB(OPC).

The improved performance of MS/G–LB(OPC) is consistently reflected not only in the solvation free energies but also in the structural properties of the first hydration shell (Figure 1 middle and bottom). In particular, the deviations in Δ*R*_1_ and Δ*n*_1_ are similar to those obtained in their respective original water models. Notably, MS/G-LB(OPC) shows improved agreement with experimental single-ion properties relative to the OPC-optimized Li-Merz force field (Figure 1, gray background). The improvement is particularly pronounced for *R*_1_ of Cl^−^ and for *n*_1_ of Cs^+^, Ca^2+^, and Br^−^.

Figure 1 also shows that it can be advantageous to select individual ion parameters that, when considered separately, outperform the MS/G-LB(OPC) combination. A particularly important example is Mg^2+^. When combined with the Mamatkulov-Schwierz Cl^−^ parameters, the microMg(OPC) model reproduces the experimental solvation free energy accurately^16^, whereas the combination with the Loche-Bonthuis Cl^−^ parameters deviates by about 10 kJ*/*mol. However, even though individual cation–anion combinations may provide higher accuracy for a particular ion many applications benefit from a consistent set of cation parameters based on a single reference anion as in MS/G-LB rather than different reference anions for each cation for instance when simulating physiological conditions with millimolar concentrations of divalent ions and 150 mM monovalent salt. The data presented here provide a reference for a large variety of ion combinations and can guide the selection of appropriate force field parameters.

### B. Activity derivative

We now assess ion–ion interactions by analyzing the activity derivative *a*_*cc*_ as an established measure for the balance between ion–ion and ion–water interactions^52^. We calculate the deviation of *a*_*cc*_ from experimental reference data for the Mamatkulov-Schwierz, Loche-Bonthuis and Herrera-Scalfi, Li-Merz, and the combined MS/G-LB(OPC) force fields (Figure 2). At the lowest investigated concentration (*c* = 0.26 m), all force fields show comparable performance. Deviations of approximately 0.1 in Δ*a*_*cc*_ can be considered good agreement within the accuracy typically achieved in force field optimization procedures.

At 1 m concentration, the monovalent cations remain well described. However, deviations are observed for Sr^2+^ with Mamatkulov-Schwierz, for Mg^2+^ and Sr^2+^ with MS/G-LB(OPC), as well as for Sr^2+^ and Ba^2+^ with the Li-Merz force field. At 2 m, the monovalent cations remain again well described. For the divalent ions, significant deviations occur for Ca^2+^, where the simulated *a*_*cc*_ is substantially underestimated for all parameter sets considered here. This behavior can be traced back to excessive ion pairing, apparent in the corresponding radial distribution functions (see SI, Fig. S7). A similar but less pronounced trend is observed for Sr^2+^. Note that for Ba^2+^ no experimental data is available at higher concentrations.

In summary, all considered parameter sets reproduce activity derivatives at low concentrations up to approximately 1 m. The monovalent cations remain well described at higher salt concentrations, whereas deviations appear for divalent ions. At higher concentrations, these deviations should be taken into consideration when comparing simulations and experiments^15^. In addition, a promising strategy to further improve the description of the divalent cations at higher concentrations is the adjustment of ion–ion combination rules beyond standard Lorentz–Berthelot mixing rules^27,28^.

### C. Kinetic properties

Finally, we provide insight into the kinetic properties of MS/G-LB(OPC) by calculating the self-diffusion coefficients and the ion-water potential of mean force (PMF). Table II summarizes the self-diffusion coefficients at infinite dilution in comparison to experimental reference data^49^. The agreement of the viscosity-scaled diffusion coefficients with experiments is comparable to that previously reported for TIP3P water^29^. However, the viscosity of OPC water is much closer to the experimental value than that of TIP3P water^41^, suggesting a more realistic dynamical description when OPC is used.

**TABLE II.** Self-diffusion coefficient *D* at infinite dilution from simulations with MS/G-LB(OPC) and experiments^49^.

| | $D^{\text{OPC}}$<br>[$10^5 \text{ cm}^2/\text{s}$] | $D^{\text{exp}}$<br>[$10^5 \text{ cm}^2/\text{s}$] |
| --- | --- | --- |
| $\text{Li}^+$ | $1.20 \pm 0.01$ | 1.029 |
| $\text{Na}^+$ | $0.93 \pm 0.02$ | 1.334 |
| $\text{K}^+$ | $1.50 \pm 0.09$ | 1.957 |
| $\text{Cs}^+$ | $1.68 \pm 0.08$ | 2.056 |
| $\text{Mg}^{2+}$ <sup>a</sup> | $0.508 \pm 0.004$ | 0.706 |
| $\text{Ca}^{2+}$ | $0.83 \pm 0.04$ | 0.792 |
| $\text{Sr}^{2+}$ | $0.76 \pm 0.03$ | 0.791 |
| $\text{Ba}^{2+}$ | $0.74 \pm 0.04$ | 0.847 |
| $\text{Cl}^-$ | $1.45 \pm 0.09$ | 2.032 |
| $\text{Br}^-$ | $1.55 \pm 0.02$ | 2.08 |

<sup>a</sup> Value for simulation taken from Ref. [16].

Figure 3 shows the ion-water PMFs, which provide first insights into water exchange between first and second hydration shell. For the monovalent cations, the free energy barrier separating the first and second minima increases with increasing charge density, as expected. Consequently, the water exchange rate with MS/G-LB(OPC) is expected to be highest for Cs^+^ and to decrease according to Cs^+^ *>* K^+^ *>* Na^+^ *>* Li^+^ in agreement with previous TIP3P results^29^ and experimental ligand exchange results^53^.

**FIG. 3.**
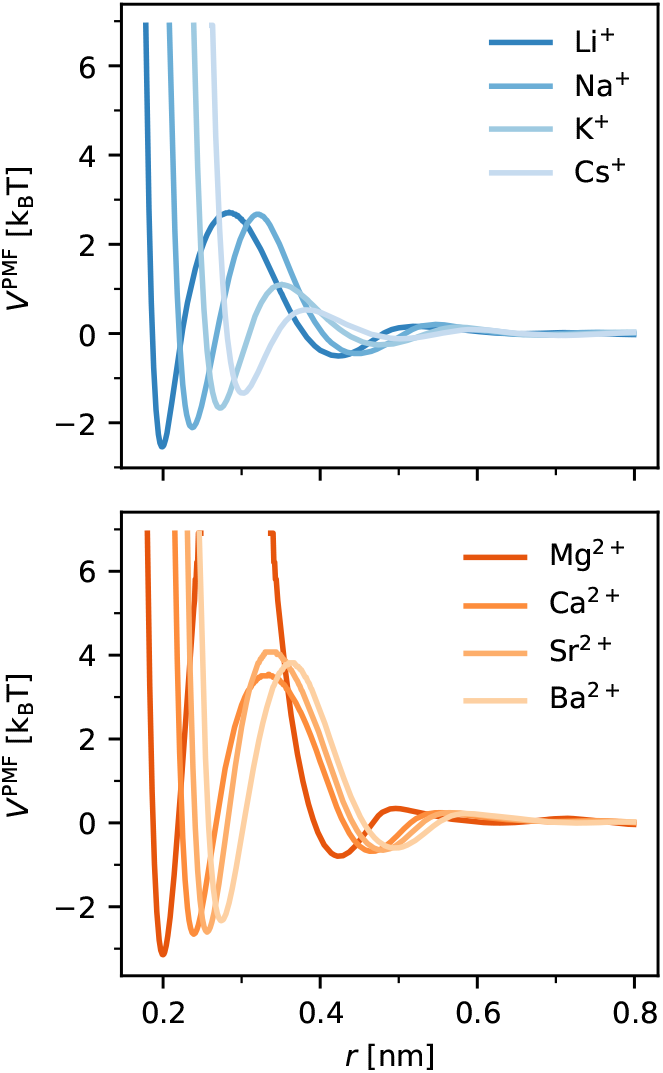
Water exchange in the first hydration shell of the cations. PMF as a function of the ion-water oxygen distance for monovalent cations (top) and divalent cations (bottom) calculated with MS/G-LB(OPC).

For the divalent cations, a similar trend is observed with the notable exception of Ca^2+^. In contrast to experimental findings, the free energy barrier for Ca^2+^ is lower than for Sr^2+^ and Ba^2+^. This indicates that the transfer of ion force fields to OPC water can lead to non-trivial and ion-specific changes in water exchange kinetics. In particular, Ca^2+^ requires additional optimization to accurately reproduce the water exchange behavior while retaining the correct single-ion and ion-pairing behavior.

## IV. CONCLUSIONS

In this work, we performed a systematic assessment of the transferability of established non-polarizable ion force fields to the four-site OPC water model for a set of monovalent (Li^+^, Na^+^, K^+^, Cs^+^) and divalent (Ca^2+^, Sr^2+^, Ba^2+^) cations together with the anions Cl^−^ and Br^−^. By benchmarking against experimental single-ion properties (hydration free energies, ion–water distances, coordination numbers) and ion-pairing properties (activity derivatives), we show that a direct transfer of existing parameter sets does not yield a uniformly accurate description of ions in OPC. Hence, we introduce the MS/G–LB(OPC) parameter combination, which reproduces hydration free energies of neutral ion pairs, first-shell structural properties and activity derivatives at low salt concentrations (up to *∼*1 m).

Our results show that transferring ion force fields to OPC can lead to unexpected and ion-specific deviations. For example, while the transfer of SPC/E-optimized parameters works well for Mg^2+^, other ions exhibit substantial discrepancies. Rather than performing a full reparameterization, we explored a limited set of Lennard–Jones parameter combinations derived from established ion models in the literature. We show that this approach already provides a viable alternative to full reparameterization: MS/G–LB(OPC) reproduces the targeted experimental observables with accuracy comparable to, and in some cases exceeding, those of the OPC-optimized Li–Merz force field. Further improvements may be achievable by exploring a broader range of Lennard–Jones parameter space beyond the subset of *σ* –*ε* values considered here. In particular, refinement of activity derivatives for divalent cations at higher salt concentrations by adjusting the ion–ion combination rules is a next natural step.

Overall, our results support the combination and transfer of ion parameters from existing force fields as a practical route for constructing ion models compatible with OPC and other water models. However, such transfers require systematic validation against experimental single-ion, ion-pairing, and kinetic properties to ensure consistent accuracy.

## V. ACKNOWLEDGEMENTS

The work was supported by the research support program (Forschungspotenziale besser nutzen!) of the University of Augsburg. The authors gratefully acknowledge the scientific support and HPC resources provided by the Erlangen National High Performance Computing Center (NHR@FAU) of the Friedrich-Alexander-Universität Erlangen-Nürnberg (FAU) under the NHR project b119ee and b253ee and the HPC resources provided on the LiCCA HPC cluster of the University of Augsburg, co-funded by the Deutsche Forschungsgemeinschaft under Project-ID 499211671.

## VI. SUPPORTING INFORMATION

Further details on the simulations and the analysis are provided in the Supporting Information: Parameters of different water models, simulation setups, details on the computation and additional results for solvation free energies, single-ion structural properties, activity derivatives, radial distribution functions and self-diffusion coefficients. The MS/G-LB(OPC) force field parameters are also available at https://git.rz.uni-augsburg.de/cbiogitpub/opc-ion-force-fields.

## VII. DATA AVAILABILITY

The MS/G-LB(OPC) force field parameters are publicly available at https://git.rz.uni-augsburg.de/cbio-gitpub/opcion-force-fields. Further data that support the finding of this study are available from the corresponding author upon reasonable request.

