## Supplementary Information for "Transferability of ion force fields to OPC water: Maintaining single-ion and ion-pairing properties"

#### Contents

**S1 Force field parameters of water models** **S3**

**S2 Simulation setup** **S3**

**S3 Calculation of solvation free energies for neutral ion pairs** **S3**

**S4 Experimental solvation free energies** **S5**

**S5 Solvation free energy for different force fields and water models** **S6**

**S6 $\Delta\Delta G_{\mathrm{solv}}$ with $Br^-$ as counterion** **S7**

**S7 Relative deviations of solvation free energies** **S7**

**S8 Solvation free energies for all anion-cation combination and selection criterion** **S8**

S8 .1 Solvation free energy of all cation-anion combinations . . . . . . . . . . . . . . . . . . . . . . . . . S8

S8 .2 Selection criterion for MS/G–LB(OPC) . . . . . . . . . . . . . . . . . . . . . . . . . . . . . S9

S8 .3 $\Delta\Delta G_{\mathrm{solv}}$ for selected parameter combinations . . . . . . . . . . . . . . . . . . . . . . . . . S9

**S9 Calculation of single-ion structural properties** **S11**

**S10 Structural properties for different force fields and water models** **S11**

**S11 Single-ion and ion-pairing properties for MS/G-LB(OPC)** **S12**

**S12 Calculation of activity derivatives** **S13**

**S13 Experimental activity derivative** **S13**

**S14 Activity derivatives for different force fields and water models** **S14**

**S15 Activity derivative for bromide salts for MS/G-LB(OPC)** **S16**

**S16 Radial distribution functions for MS/G-LB(OPC)** **S17**

**S17 Calculation of self-diffusion coefficients** **S19**

**S18 Self-diffusion coefficient for different force fields and water models** **S20**

**S19 Bibliography** **S21**

### S1 Force field parameters of water models

Table S1: Parameters of water models relevant for this work taken from Ref. 1. For the 3-site water models TIP3P[2] and SPC/E,[3] the negative partial charge ($q_{\mathrm{Ow/Mw}}$) is located on the oxygen Ow. For the 4-site water model OPC,[1] it is placed on the dummy atom Mw. $q_{\mathrm{H}}$ is the partial charge of the hydrogen atoms and $\sigma_{\mathrm{Ow}}$ and $\varepsilon_{\mathrm{Ow}}$ are the Lennard-Jones parameters of the water oxygen atom. The length of the oxygen hydrogen bond is $l$, the angle between the two bonds in the water molecule is $\theta$. The distance between Ow and Mw is given for OPC as $z$.

| | $q_{\mathrm{Ow/Mw}}$ [e] | $q_{\mathrm{H}}$ [e] | $\sigma_{\mathrm{Ow}}$ [nm] | $\varepsilon_{\mathrm{Ow}}$ [kJ/mol] | $l$ [nm] | $\theta$ [°] | $z$ [nm] |
|---|---|---|---|---|---|---|---|
| TIP3P | -0.834 | 0.417 | 0.315061 | 0.6364 | 0.09572 | 104.52 | n.a. |
| SPC/E | -0.8476 | 0.4238 | 0.3166 | 0.650 | 0.1 | 109.47 | n.a. |
| **OPC** | **-1.3582** | **0.6791** | **0.316655** | **0.89036** | **0.08724** | **103.6** | **0.01594** |
| exp. (gas) | n.a. | n.a. | n.a. | n.a. | 0.09572 | 104.54 | n.a |

### S2 Simulation setup

Table S2: Simulation setups. 'Phys. property' indicates the physical property that was obtained from the corresponding type of simulation using either free energy perturbation (FEP) or unbiased simulations as 'Method' . 'System' lists all particles of this respective simulation type and 'Duration' the simulation time (the products indicate the number of windows times simulation time in case of FEP or the number of independent simulations times simulation time in case of simulations for $a_{cc}$ or $D$). '$L$' denotes the simulation box size. In all cases a cubic box was employed. 'Ensemble' expresses if a canonical ensemble (NVT), or an isobaric-isothermal ensemble (NPT) was used. We here use $Ca^{2+}$ and $Cl^{-}$ as placeholders, equivalent simulations were run for each cation and anion combination.

| Phys. property | Method | System | Duration | $L$ | Ensemble |
|---|---|---|---|---|---|
| $R_1$, $n_1$, $\Delta G_{\mathrm{solv}}$ | FEP | 1 ion, 506 water | 40 × 1 ns | 2.5 nm | NPT |
| $D$ | unbiased | 1 ion, 506 water | 3 × 50 ns | 2.5 nm | NVT |
| $a_{\mathrm{cc}}$ | unbiased | 10 $Li^{+}$, 10 $Cl^{-}$, 2171 water (0.26 m) | 3 × 150 ns | 4 nm | NPT |
| $a_{\mathrm{cc}}$ | unbiased | 20 $Li^{+}$, 20 $Cl^{-}$, 2135 water (0.52 m) | 3 × 150 ns | 4 nm | NPT |
| $a_{\mathrm{cc}}$ | unbiased | 39 $Li^{+}$, 39 $Cl^{-}$, 2048 water (1.06 m) | 3 × 150 ns | 4 nm | NPT |
| $a_{\mathrm{cc}}$ | unbiased | 73 $Li^{+}$, 73 $Cl^{-}$, 1909 water (2.12 m) | 3 × 150 ns | 4 nm | NPT |
| $a_{\mathrm{cc}}$ | unbiased | 10 $Ca^{2+}$, 20 $Cl^{-}$, 2171 water (0.26 m) | 3 × 150 ns | 4 nm | NPT |
| $a_{\mathrm{cc}}$ | unbiased | 20 $Ca^{2+}$, 40 $Cl^{-}$, 2135 water (0.52 m) | 3 × 150 ns | 4 nm | NPT |
| $a_{\mathrm{cc}}$ | unbiased | 39 $Ca^{2+}$, 78 $Cl^{-}$, 2048 water (1.06 m) | 3 × 150 ns | 4 nm | NPT |
| $a_{\mathrm{cc}}$ | unbiased | 73 $Ca^{2+}$, 146 $Cl^{-}$, 1909 water (2.12 m) | 3 × 150 ns | 4 nm | NPT |
| $V_{\mathrm{PMF}}$ | unbiased | 39 $Li^{+}$, 39 $Cl^{-}$, 2048 water (1.06 m) | 1000 ns | 4 nm | NPT |
| $V_{\mathrm{PMF}}$ | unbiased | 39 $Ca^{2+}$, 78 $Cl^{-}$, 2048 water (1.06 m) | 1000 ns | 4 nm | NPT |

### S3 Calculation of solvation free energies for neutral ion pairs

The calculation of solvation free energies has been described in several previous works such as.[4–6]

Briefly, free energy perturbation (FEP) simulations in combination with Bennet's acceptance ratio (BAR) method[7] were employed to determine the solvation free energies of neutral ion pairs. The FEP simulations were performed for each ion parameterization over 40 evenly spaced intermediate $\lambda$-windows. Each window was simulated for 1 ns. The first 20 windows created a neutral Lennard-Jones particle and during the second 20 windows the magnitude of the charge was increased until it reached the desired value, *i.e.* -1 for anions, +1 for monovalent cations, and +2 for divalent cations. Soft-core potentials were employed to avoid divergences. For our analyses, we excluded the first 200 ps from each window for equilibration.

In order to directly compare with experiments, we applied several corrections and only report solvation free energies for charge neutral ion pairs. Finite size effects were taken into account by:[8]

$$\Delta G_{\mathrm{fs}} = \frac{z^2 N_A e^2}{4\pi\epsilon_0}\left[-\frac{\zeta_{\mathrm{ew}}}{2\epsilon_r} + \left(1-\frac{1}{\epsilon_r}\right)\left(\frac{2\pi R_1^2}{3L^3} - \frac{8\pi^2 R_1^5}{45L^6}\right)\right] , \tag{S1}$$

In this equation, $z$ denotes the valency, $N_A$ Avogadro's number, $e$ is the elementary charge, $\epsilon_0$ the vacuum permittivitty, $R_1$ is the first peak of the ion-water radial distribution function and $\zeta_{ew} = -2.837\,297/L$ is the Wigner potential for the edge length $L$ of a cubic simulation box.[8,9] $\epsilon_r$ is the relative dielectric constant of the different water models and we employed 83 for TIP3P, 71 for SPC/E and 78.4 for OPC.[1,5,10]

The compression of an ideal gas with $p_0 \approx 1$ bar to the pressure of an ideal solution at a density of 1 mol/l with $p_1 = 26.4$ bar can be accounted for with the following correction term:[4]

$$\Delta G_{\mathrm{press}} = N_{\mathrm{A}} k_{\mathrm{B}} T \ln(p_1/p_0) = 7.9 \text{ kJ/mol} . \tag{S2}$$

For comparison with experimental data, we exclusively consider *neutral ion pairs.* Experimental solvation free energies are more robust for such pairs because they do not depend on the proton solvation free energy, which is known to be subject to significant uncertainty.[11] In addition, for neutral ion pairs the interfacial crossing term cancels, eliminating contributions from the water-model–dependent surface potential, which can otherwise introduce systematic errors. Here, we use halide counterions ($Cl^-$ or $Br^-$) avoiding the need to reference the proton solvation free energy and the surface-potential contribution, making comparisons between simulation and experiment more reliable.

The solvation free energy of the neutral salt pair is obtained as the sum of the cation and chloride or bromide contributions:

$$\Delta G_{\mathrm{solv}} := \Sigma_{\Delta G} = \Delta G_{\mathrm{solv}}^{\mathrm{cation}} + z \times \Delta G_{\mathrm{solv}}^{\mathrm{anion}}, \tag{S3}$$

where $z$ is the charge of the cation.

For completeness, Table S16 reports the simulated solvation free energies of the individual ions (cations and anions). These values are provided as a reference for future simulation studies and are *not* used for comparison with experimental data. In this case, the interfacial crossing term was explicitly included in the calculation and was calculated as[4]

$$\Delta G_{\text{surf}} = N_A z \cdot e\phi_{\text{surf}} = -z \cdot 50.8 \text{ kJ/mol} , \tag{S4}$$

with a surface potential of $\phi_{\text{surf}} = -0.527\,\text{V}$,[12,13] a value that closely matches the experimental estimation of $-0.50\,V$.[11] The simulated solvation free energy of a single cation is obtained as:

$$\Delta G^{\text{cation}}_{\text{solv}} = \Delta G^{\text{cation}}_{\text{sim}} + \Delta G^{\text{cation}}_{\text{fs}} + \Delta G^{\text{cation}}_{\text{surf}} + \Delta G^{\text{cation}}_{\text{press}} . \tag{S5}$$

### S4 Experimental solvation free energies

All simulations were performed at 300 K to remain consistent with previous force field studies conducted at this temperature. Experimental solvation free energies, however, are typically reported at 298.15 K. To enable a direct comparison, we converted the experimental values to 300 K. Following the procedure of Loche *et al.*,[14] the solvation free energy at 300 K was estimated from the reported experimental enthalpy and entropy of solvation using

$$\Delta G(T) = \Delta H - T\Delta S. \tag{S6}$$

We evaluated $\Delta G$ at $T = 300\,\text{K}$ using the experimental values of $\Delta H$ and $\Delta S$. In addition, we take average values of $\Delta G$ reported by Marcus[15] and Tissandier[16] for the monovalent cations $Li^+$, Na,$^+$ and $K^+$. In case of $Cs^+$ and the divalent cations $Ca^{2+}$, $Sr^{2+}$ and $Ba^{2+}$, the corresponding data is only reported by Marcus,[15] and therefore no averaging was performed.

#### S5 Solvation free energy for different force fields and water models

Table S3: Solvation free energies of neutral ion pairs $\Delta G_{\mathrm{solv}}$ in kJ/mol with $Cl^-$ as counterion. Simulation results are shown for the following force-field sets: Mamatkulov–Schwierz[5] (MS), Fyta–Netz/Mamatkulov-Netz[17,18] (FN/MN), Loche–Bonthuis and Herrera-Scalfi parameters[14,19] (LB/HS), and Li–Merz hydration free energy (HFE) parameter set[20,21] (LM). In addition, results for the combination of cation parameters from MS, except for Mg, for which the microMg parameters for OPC water[22] were employed, and anion parameters from LB/HS (MS/G-LB) as well as experimental reference values (Exp.) are shown. The current MS/G-LB(OPC) is highlighted.

| | $Li^+$ | $Na^+$ | $K^+$ | $Cs^+$ | $Mg^{2+}$ | $Ca^{2+}$ | $Sr^{2+}$ | $Ba^{2+}$ |
|---|---|---|---|---|---|---|---|---|
| MS (TIP3P) | -826.5 | -721.2 | -648.7 | -603.1 | -2530.2 | -2209.4 | -2079.2 | -1951.4 |
| MS (OPC) | -826.2 | -723.2 | -650.7 | -604.5 | -2512.3 | -2198.7 | -2071.2 | -1943.0 |
| FN/MN (SPC/E) | – | -690.0 | -653.5 | -608.1 | -2533.3 | -2206.4 | -2077.2 | -1953.3 |
| FN/MN (OPC) | – | -692.3 | -657.0 | -608.9 | -2497.3 | -2185.3 | -2060.4 | -1939.9 |
| LB/HS (SPC/E) | -839.2 | -730.5 | -661.0 | -614.2 | – | – | – | – |
| LB/HS (OPC) | -837.8 | -734.1 | -664.3 | -617.1 | – | – | – | – |
| LM (OPC) | -823.9 | -709.6 | -635.9 | -590.9 | -2537.6 | -2208.4 | -2079.9 | -1949.6 |
| **MS/G-LB(OPC)** | -830.0 | -727.0 | -654.5 | -608.3 | -2548.6 | -2206.3 | -2078.8 | -1950.6 |
| Exp. | -833.0 | -727.4 | -656.1 | -609.8 | 2538.7 | -2215.4 | -2086.4 | -1959.5 |

Table S4: Solvation free energies of neutral ion pairs $\Delta G_{\mathrm{solv}}$ in kJ/mol with $Br^-$ as counterion. The current MS/G-LB(OPC) is highlighted.

| | $Li^+$ | $Na^+$ | $K^+$ | $Cs^+$ | $Mg^{2+}$ | $Ca^{2+}$ | $Sr^{2+}$ | $Ba^{2+}$ |
|---|---|---|---|---|---|---|---|---|
| FN/MN (SPC/E) | – | -667.2 | -630.7 | -585.3 | -2487.8 | -2160.8 | -2031.6 | -1907.7 |
| FN/MN (OPC) | – | -669.5 | -634.3 | -586.2 | -2451.8 | -2139.8 | -2014.9 | -1894.4 |
| LB/HS (SPC/E) | -814.8 | -706.0 | -636.5 | -589.7 | – | – | – | – |
| LB/HS (OPC) | -813.8 | -710.1 | -640.3 | -593.1 | – | – | – | – |
| LM (OPC) | -802.2 | -687.9 | -614.2 | -569.2 | -2494.2 | -2165.0 | -2036.5 | -1906.2 |
| **MS/G-LB(OPC)** | -806.0 | -703.0 | -630.5 | -584.3 | -2500.6 | -2158.3 | -2030.9 | -1902.6 |
| Exp | -806.6 | -700.9 | -629.6 | -583.6 | -2486.3 | -2163.0 | -2034.0 | -1907.1 |

#### S6 $\Delta\Delta G_{\mathrm{solv}}$ with $Br^-$ as counterion

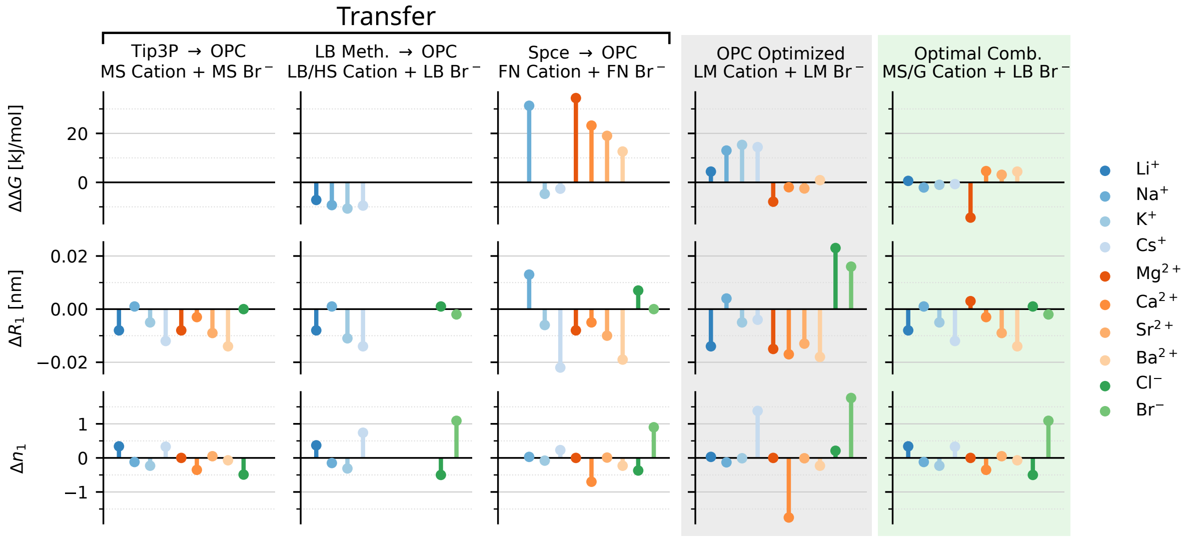


Figure S1: **Difference between simulated and experimental single-ion properties:** $\Delta\Delta G_{\mathrm{solv}}$ with $Br^-$ as counterion (top), $\Delta R_1$ (middle) and $\Delta n_1$ (bottom). Results for transferring three not-OPC optimized parameter sets to OPC (white background). Results for the OPC-optimized Li-Merz parameters (gray background). Optimal combination of cation and anion set in OPC (green background). The experimental values are from Refs. 15,16,23.

#### S7 Relative deviations of solvation free energies

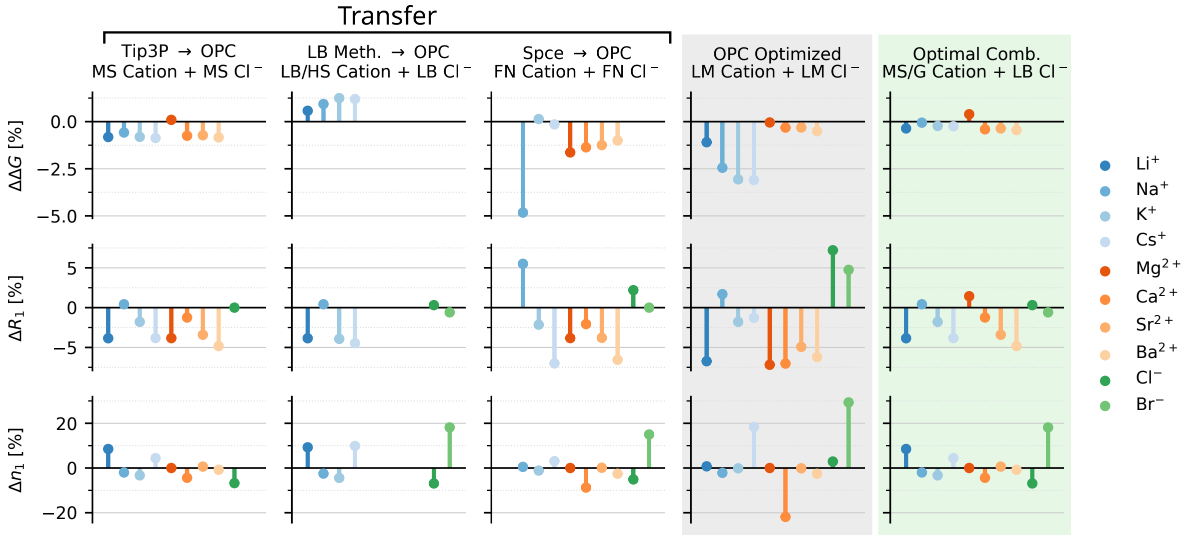


Figure S2: **Relative error of simulated single-ion properties with respect to their experimental reference:** $\Delta\Delta G_{\mathrm{solv}}$ with $Cl^-$ as counterion (top), $\Delta R_1$ (middle) and $\Delta n_1$ (bottom). Results for transferring three not-OPC optimized parameter sets to OPC (white background). Results for the OPC-optimized Li-Merz parameters (gray background). Optimal combination of cation and anion set in OPC (green background). The experimental values are from Refs. 15,16,23.

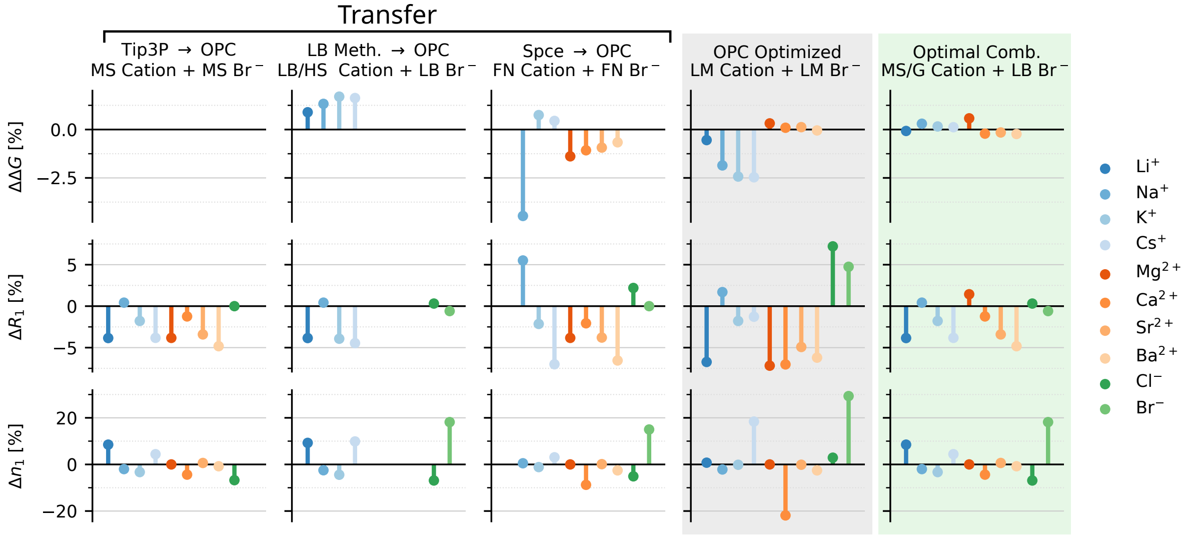


Figure S3: **Relative error of simulated single-ion properties with respect to their experimental reference:** $\Delta\Delta G_{\mathrm{solv}}$ with $\mathrm{Br}^-$ as counterion (top), $\Delta R_1$ (middle) and $\Delta n_1$ (bottom). Results for transferring three not-OPC optimized parameter sets to OPC (white background). Results for the OPC-optimized Li-Merz parameters (gray background). Optimal combination of cation and anion set in OPC (green background). The experimental values are from Refs. 15,16,23.

### S8 Solvation free energies for all anion-cation combination and selection criterion

#### S8 .1 Solvation free energy of all cation-anion combinations

The solvation free energy of all cation-anion combinations in OPC water can be calculated from:

$$\Delta G_{\mathrm{solv}} = \Sigma_{\Delta G} = \Delta G_{\mathrm{solv}}^{\mathrm{cation}} + z \times \Delta G_{\mathrm{solv}}^{\mathrm{anion}}, \tag{S7}$$

Here $\Delta G_{\mathrm{solv}}^{\mathrm{cation}}$ and $\Delta G_{\mathrm{solv}}^{\mathrm{anion}}$ are the simulated solvation free energies of individual ions with applied finite size and pressure correction (see section S3). The values are listed in Table S5.

Table S5: Simulated solvation free energies of anions and cations ($\Delta G_{\mathrm{solv}}^{\mathrm{cation}}$ and $\Delta G_{\mathrm{solv}}^{\mathrm{anion}}$) with applied finite size and pressure correction in kJ/mol.

| | $Li^+$ | $Na^+$ | $K^+$ | $Cs^+$ | $Mg^{2+}$ | $Ca^{2+}$ | $Sr^{2+}$ | $Ba^{2+}$ | $Cl^-$ | $Br^-$ |
|---|---|---|---|---|---|---|---|---|---|---|
| MS/G | -508.8 | -405.8 | -333.4 | -287.1 | -1906.3 | -1564.0 | -1436.5 | -1308.3 | -317.4 | – |
| LB/HS | -516.6 | -413.0 | -343.1 | -295.9 | – | – | – | – | -321.2 | -297.2 |
| FN/MN | – | -381.8 | -346.5 | -298.4 | -1876.3 | -1564.2 | -1439.4 | -1318.9 | -310.5 | -287.8 |
| LM | -517.7 | -403.4 | -329.8 | -284.7 | -1925.2 | -1596.0 | -1467.5 | -1337.2 | -306.2 | -284.5 |

#### S8 .2 Selection criterion for MS/G–LB(OPC)

For the selection of the MS/G–LB(OPC) force field parameters, we consider the solvation free energy as the primary target property. Among the parameter sets examined, MS/G–LB(OPC) provides the smallest overall deviation across a complete set of eight cations when paired with a fixed reference anion. The corresponding deviations for the cation sets with a fixed reference anion are listed below. We note that a few individual cation–anion combinations (e.g., FN Cs/FN Cl or LM Mg/LM Cl) yield slightly higher accuracy for these specific ions in OPC water. However, many applications benefit from a transferable and internally consistent set based on a single reference anion. From this perspective, MS/G–LB(OPC) represents the most balanced choice, providing the best overall agreement with solvation free energies as well as structural properties and ion-pairing.

#### S8 .3 $\Delta\Delta G_{\text{solv}}$ for selected parameter combinations

Tables S6-S13 show the deviations of the solvation free energies of neutral ion pairs from experiments for selected parameter combinations. Among the parameter combinations covering the complete set of eight cations and both anions, MS/G–LB yields the smallest overall deviation from experimental solvation free energies (mean absolute deviation 4.6 kJ mol$^{-1}$). The agreement is particularly good for monovalent salts (mean absolute deviation 1.3 kJ mol$^{-1}$), while larger deviations are observed for divalent salts (mean absolute deviation 7.8 kJ mol$^{-1}$).

Table S6: Deviations of solvation free energies of neutral ion pairs from experimental reference values, $\Delta\Delta G_{\text{solv}} = \Delta G_{\text{solv}}^{\text{sim}} - \Delta G_{\text{solv}}^{\text{exp}}$, in kJ/mol for combinations of MS/G cation parameters with the four parameterizations for $Cl^-$.

| | $Li^+$ | $Na^+$ | $K^+$ | $Cs^+$ | $Mg^{2+}$ | $Ca^{2+}$ | $Sr^{2+}$ | $Ba^{2+}$ |
|---|---|---|---|---|---|---|---|---|
| $Cl^-$ (MS) | 6.8 | 4.2 | 5.4 | 5.3 | -2.3 | 16.7 | 15.2 | 16.5 |
| $Cl^-$ (LB/HS) | 3.0 | 0.4 | 1.6 | 1.5 | -9.9 | 9.1 | 7.6 | 8.9 |
| $Cl^-$ (FN/MN) | 13.7 | 11.1 | 12.2 | 12.1 | 11.4 | 30.4 | 28.8 | 30.2 |
| $Cl^-$ (LM) | 18.0 | 15.4 | 16.6 | 16.5 | 20.1 | 39.1 | 37.5 | 38.9 |

Table S7: Deviations of solvation free energies of neutral ion pairs from experimental reference values, $\Delta\Delta G_{\text{solv}} = \Delta G_{\text{solv}}^{\text{sim}} - \Delta G_{\text{solv}}^{\text{exp}}$, in kJ/mol for combinations of MS/G cation parameters with the three parameterizations for $Br^-$.

| | $Li^+$ | $Na^+$ | $K^+$ | $Cs^+$ | $Mg^{2+}$ | $Ca^{2+}$ | $Sr^{2+}$ | $Ba^{2+}$ |
|---|---|---|---|---|---|---|---|---|
| $Br^-$ (LB/HS) | 0.5 | -2.1 | -0.9 | -0.7 | -14.3 | 4.7 | 3.1 | 4.5 |
| $Br^-$ (FN/MN) | 9.9 | 7.3 | 8.5 | 8.7 | 4.5 | 23.5 | 21.9 | 23.3 |
| $Br^-$ (LM) | 13.2 | 10.6 | 11.8 | 12.0 | 11.1 | 30.0 | 28.5 | 29.9 |

Table S8: Deviations of solvation free energies of neutral ion pairs from experimental reference values, $\Delta\Delta G_{\text{solv}} = \Delta G_{\text{solv}}^{\text{sim}} - \Delta G_{\text{solv}}^{\text{exp}}$, in kJ/mol for combinations of LB/HS cation parameters with the four parameterizations for $Cl^-$.

| | $Li^+$ | $Na^+$ | $K^+$ | $Cs^+$ |
|---|---|---|---|---|
| $Cl^-$ (MS) | -0.9 | -2.9 | -4.4 | -3.5 |
| $Cl^-$ (LB/HS) | -4.7 | -6.7 | -8.1 | -7.3 |
| $Cl^-$ (FN/MN) | 5.9 | 4.0 | 2.5 | 3.3 |
| $Cl^-$ (LM) | 10.2 | 8.3 | 6.8 | 7.7 |

Table S9: Deviations of solvation free energies of neutral ion pairs from experimental reference values, $\Delta\Delta G_{\text{solv}} = \Delta G_{\text{solv}}^{\text{sim}} - \Delta G_{\text{solv}}^{\text{exp}}$, in kJ/mol for combinations of LB/HS cation parameters with the three parameterizations for $Br^-$.

| | $Li^+$ | $Na^+$ | $K^+$ | $Cs^+$ |
|---|---|---|---|---|
| $Br^-$ (LB/HS) | -7.2 | -9.2 | -10.6 | -9.5 |
| $Br^-$ (FN/MN) | 2.2 | 0.2 | -1.2 | -0.1 |
| $Br^-$ (LM) | 5.5 | 3.5 | 2.0 | 3.2 |

Table S10: Deviations of solvation free energies of neutral ion pairs from experimental reference values, $\Delta\Delta G_{\text{solv}} = \Delta G_{\text{solv}}^{\text{sim}} - \Delta G_{\text{solv}}^{\text{exp}}$, in kJ/mol for combinations of FN/MN cation parameters with the four parameterizations for $Cl^-$.

| | $Na^+$ | $K^+$ | $Cs^+$ | $Mg^{2+}$ | $Ca^{2+}$ | $Sr^{2+}$ | $Ba^{2+}$ |
|---|---|---|---|---|---|---|---|
| $Cl^-$ (MS) | 28.3 | -7.7 | -6.0 | 27.7 | 16.4 | 12.3 | 5.9 |
| $Cl^-$ (LB/HS) | 24.5 | -11.5 | -9.8 | 20.1 | 8.9 | 4.7 | -1.7 |
| $Cl^-$ (FN/MN) | 35.1 | -0.9 | 0.9 | 41.4 | 30.1 | 26.0 | 19.6 |
| $Cl^-$ (LM) | 39.5 | 3.4 | 5.2 | 50.0 | 38.8 | 34.7 | 28.2 |

Table S11: Deviations of solvation free energies of neutral ion pairs from experimental reference values, $\Delta\Delta G_{\text{solv}} = \Delta G_{\text{solv}}^{\text{sim}} - \Delta G_{\text{solv}}^{\text{exp}}$, in kJ/mol for combinations of FN/MN cation parameters with the three parameterizations for $Br^-$.

| | $Na^+$ | $K^+$ | $Cs^+$ | $Mg^{2+}$ | $Ca^{2+}$ | $Sr^{2+}$ | $Ba^{2+}$ |
|---|---|---|---|---|---|---|---|
| $Br^-$ (LB/HS) | 22.0 | -14.0 | -12.0 | 15.7 | 4.4 | 0.3 | -6.1 |
| $Br^-$ (FN/MN) | 31.4 | -4.6 | -2.6 | 34.5 | 23.2 | 19.1 | 12.7 |
| $Br^-$ (LM) | 34.7 | -1.3 | 0.7 | 41.0 | 29.8 | 25.6 | 19.2 |

Table S12: Deviations of solvation free energies of neutral ion pairs from experimental reference values, $\Delta\Delta G_{\text{solv}} = \Delta G_{\text{solv}}^{\text{sim}} - \Delta G_{\text{solv}}^{\text{exp}}$, in kJ/mol for combinations of LM cation parameters with the four parameterizations for $Cl^-$.

| | $Li^+$ | $Na^+$ | $K^+$ | $Cs^+$ | $Mg^{2+}$ | $Ca^{2+}$ | $Sr^{2+}$ | $Ba^{2+}$ |
|---|---|---|---|---|---|---|---|---|
| $Cl^-$ (MS) | -2.0 | 6.7 | 9.0 | 7.7 | -21.2 | -15.3 | -15.8 | -12.4 |
| $Cl^-$ (LB/HS) | -5.8 | 2.9 | 5.2 | 3.9 | -28.8 | -22.9 | -23.4 | -20.0 |
| $Cl^-$ (FN/MN) | 4.8 | 13.5 | 15.8 | 14.6 | -7.6 | -1.6 | -2.1 | 1.2 |
| $Cl^-$ (LM) | 9.1 | 17.9 | 20.2 | 18.9 | 1.1 | 7.0 | 6.5 | 9.9 |

Table S13: Deviations of solvation free energies of neutral ion pairs from experimental reference values, $\Delta\Delta G_{\mathrm{solv}} = \Delta G^{\mathrm{sim}}_{\mathrm{solv}} - \Delta G^{\mathrm{exp}}_{\mathrm{solv}}$, in kJ/mol for combinations of LM cation parameters with the three parameterizations for $Br^-$.

| | $Li^+$ | $Na^+$ | $K^+$ | $Cs^+$ | $Mg^{2+}$ | $Ca^{2+}$ | $Sr^{2+}$ | $Ba^{2+}$ |
|---|---|---|---|---|---|---|---|---|
| $Br^-$ (LB) | -8.3 | 0.4 | 2.7 | 1.7 | -33.3 | -27.3 | -27.8 | -24.5 |
| $Br^-$ (FN/MN) | 1.1 | 9.8 | 12.1 | 11.1 | -14.5 | -8.5 | -9.0 | -5.7 |
| $Br^-$ (LM) | 4.4 | 13.1 | 15.4 | 14.4 | -7.9 | -2.0 | -2.5 | 0.9 |

### S9 Calculation of single-ion structural properties

Single-ion structural properties were calculated in the same way as in our previous work.[6] For this, the radial distribution functions $g(r)$ between the ions and water oxygen atoms were employed. The first-shell radius $R_1$ was obtained as the position of the first maximum of $g(r)$. The average coordination number $n_1$ was determined by integrating $g(r)$:

$$n_1 = 4\pi\rho \int_0^{r_{\mathrm{min}}} g(r) r^2 \mathrm{d}r, \tag{S8}$$

where $\rho$ is the bulk water density and $r_{\mathrm{min}}$ is the first minimum of $g(r)$. The $g(r)$ functions were computed from a 1 ns NPT trajectory of the final FEP state containing one ion and 506 water molecules. The uncertainty in $R_1$ is approximately $\pm 0.004$ nm. Experimental reference values were taken from Ref. 23. For $n_1$, the number in parentheses denotes the experimental value used for comparison.

### S10 Structural properties for different force fields and water models

Table S14: Ion–oxygen distance of the first hydration shell $R_1$ (nm) for ions in water. Experimental values are taken from Ref. 23. The current MS/G-LB(OPC) is highlighted.

| | $Li^+$ | $Na^+$ | $K^+$ | $Cs^+$ | $Mg^{2+}$ | $Ca^{2+}$ |
|---|---|---|---|---|---|---|
| MS (TIP3P) | 0.196 | 0.233 | 0.269 | 0.297 | 0.195 | 0.237 |
| **MS/G (OPC)** | 0.200 | 0.237 | 0.275 | 0.302 | 0.212 | 0.239 |
| FN/MN (SPC/E) | – | 0.247 | 0.269 | 0.288 | 0.196 | 0.232 |
| FN/MN (OPC) | – | 0.249 | 0.274 | 0.292 | 0.201 | 0.237 |
| LB/HS (SPC/E) | 0.194 | 0.235 | 0.267 | 0.298 | – | – |
| **LB/HS (OPC)** | 0.200 | 0.237 | 0.269 | 0.300 | – | – |
| LM | 0.194 | 0.240 | 0.275 | 0.310 | 0.194 | 0.225 |
| Exp | 0.208 ± 0.006 | 0.236 ± 0.006 | 0.280 ± 0.008 | 0.314 ± 0.006 | 0.209 ± 0.004 | 0.242 ± 0.005 |

| | $Sr^{2+}$ | $Ba^{2+}$ | $Cl^-$ | $Br^-$ |
|---|---|---|---|---|
| MS (TIP3P) | 0.254 | 0.273 | 0.319 | – |
| **MS/G (OPC)** | 0.255 | 0.276 | 0.319 | – |
| FN/MN (SPC/E) | 0.250 | 0.266 | 0.323 | 0.334 |
| FN/MN (OPC) | 0.254 | 0.271 | 0.326 | 0.337 |
| LB/HS (SPC/E) | – | – | 0.317 | 0.332 |
| **LB/HS (OPC)** | – | – | 0.320 | 0.335 |
| LM | 0.251 | 0.272 | 0.342 | 0.354 |
| Exp | 0.264 ± 0.004 | 0.290 ± 0.006 | 0.319 ± 0.007 | 0.337 ± 0.006 |

Table S15: Coordination number $n_1$ for ions in water. Experimental values are taken from Ref. 23. The current MS/G-LB(OPC) is highlighted.

| | $Li^+$ | $Na^+$ | $K^+$ | $Cs^+$ | $Mg^{2+}$ | $Ca^{2+}$ | $Sr^{2+}$ | $Ba^{2+}$ | $Cl^-$ | $Br^-$ |
|---|---|---|---|---|---|---|---|---|---|---|
| MS (TIP3P) | 4.26 | 5.70 | 6.78 | 7.99 | 6 | 7.82 | 8.13 | 8.91 | 7.62 | – |
| **MS/G (OPC)** | 4.34 | 5.88 | 6.77 | 7.83 | 6 | 7.65 | 8.05 | 8.93 | 6.76 | – |
| FN/MN (SPC/E) | – | 5.95 | 6.99 | 7.63 | 6 | 7.22 | 8.00 | 8.44 | 7.28 | 7.32 |
| FN/MN (OPC) | – | 6.03 | 6.92 | 7.73 | 6 | 7.30 | 8.01 | 8.77 | 6.88 | 6.90 |
| LB/HS (SPC/E) | 4.16 | 5.68 | 6.62 | 8.26 | – | – | – | – | 7.20 | 7.43 |
| **LB/HS (OPC)** | 4.37 | 5.85 | 6.69 | 8.24 | – | – | – | – | 6.75 | 6.83 |
| LM | 4.03 | 5.87 | 6.99 | 8.88 | 6 | 6.25 | 7.99 | 8.77 | 7.46 | 8.17 |
| Exp | 4–6 (4) | 4–8 (6) | 6–8 | 7–8 | 6 | 8 | 7.9–8 | 9 | 6–8.5 | 6 |

### S11 Single-ion and ion-pairing properties for MS/G-LB(OPC)

Table S16: **Results for single-ion properties in OPC water obtained with the MS/G-LB(OPC) force field parameters introduced in this work.** Shown are the solvation free energy for single cations $\Delta G_{\mathrm{solv}}^{\mathrm{cation}}$/anions $\Delta G_{\mathrm{solv}}^{\mathrm{anion}}$, the coordination number of the first hydration shell $n_1$, and the ion-oxygen distance in the first hydration shell $R_1$.

| Ion | $\Delta G_{\mathrm{solv}}^{\mathrm{cation/anion}}$ (kJ/mol) | $n_1$ | $n_1^{\mathrm{exp}}$ [23] | $R_1$ (nm) | $R_1^{\mathrm{exp}}$ (nm) [23] |
|---|---|---|---|---|---|
| $Li^+$ | -508.8 ± 1 | 4.34 | 4–6 (4) | 0.200 ± 0.004 | 0.208 ± 0.006 |
| $Na^+$ | -405.8 ± 1 | 5.88 | 4–8 (6) | 0.237 ± 0.004 | 0.236 ± 0.006 |
| $K^+$ | -333.4 ± 1 | 6.77 | 6–8 | 0.275 ± 0.004 | 0.280 ± 0.008 |
| $Cs^+$ | -287.1 ± 1 | 7.83 | 7–8 | 0.302 ± 0.004 | 0.314 ± 0.006 |
| $Mg^{2+}$ | -1906.3 ± 1 | 6 [22] | 6 | 0.212 ± 0.004 [22] | 0.209 ± 0.004 |
| $Ca^{2+}$ | -1564.0 ± 1 | 7.65 | 8 | 0.239 ± 0.004 | 0.242 ± 0.005 |
| $Sr^{2+}$ | -1436.5 ± 1 | 8.05 | 7.9–8 | 0.255 ± 0.004 | 0.264 ± 0.004 |
| $Ba^{2+}$ | -1308.3 ± 1 | 8.93 | 9 | 0.276 ± 0.004 | 0.290 ± 0.006 |
| $Cl^-$ | -321.2 ± 1 | 6.75 | 6–8.5 | 0.320 ± 0.004 | 0.319 ± 0.007 |
| $Br^-$ | -297.2 ± 1 | 6.83 | 6 | 0.335 ± 0.004 | 0.337 ± 0.006 |

Table S17: **Results for ion-parining properties in OPC water obtained with the MS/G-LB(OPC) force field parameters introduced in this work.** Activity derivatives $a_{cc}$ in OPC water for chloride salts from MS/G-LB(OPC) at four concentrations: 0.26, 0.52, 1.06, and 2.12 m.

| | 0.26 m | | 0.52 m | | 1.06 m | | 2.12 m | |
|---|---|---|---|---|---|---|---|---|
| Ion | $a_{cc}^{\mathrm{sim}}$ | $a_{cc}^{\mathrm{exp}}$ | $a_{cc}^{\mathrm{sim}}$ | $a_{cc}^{\mathrm{exp}}$ | $a_{cc}^{\mathrm{sim}}$ | $a_{cc}^{\mathrm{exp}}$ | $a_{cc}^{\mathrm{sim}}$ | $a_{cc}^{\mathrm{exp}}$ |
| $Li^+$ | $0.95 \pm 0.04$ | 0.96 | $1.02 \pm 0.03$ | 1.01 | $1.07 \pm 0.01$ | 1.15 | $1.30 \pm 0.03$ | 1.47 |
| $Na^+$ | $0.99 \pm 0.01$ | 0.92 | $0.97 \pm 0.02$ | 0.93 | $1.04 \pm 0.03$ | 0.97 | $1.12 \pm 0.01$ | 1.12 |
| $K^+$ | $0.92 \pm 0.02$ | 0.89 | $0.94 \pm 0.02$ | 0.89 | $0.94 \pm 0.01$ | 0.90 | $0.97 \pm 0.02$ | 0.96 |
| $Cs^+$ | $0.88 \pm 0.02$ | 0.86 | $0.87 \pm 0.02$ | 0.84 | $0.92 \pm 0.01$ | 0.84 | $0.96 \pm 0.03$ | 0.89 |
| $Mg^{2+}$ | $0.84 \pm 0.02$ | 0.86 | $0.96 \pm 0.02$ | 0.93 | $1.31 \pm 0.04$ | 1.53 | $2.08 \pm 0.11$ | 2.67 |
| $Ca^{2+}$ | $0.91 \pm 0.01$ | 0.90 | $1.10 \pm 0.02$ | 1.03 | $1.50 \pm 0.07$ | 1.38 | $0.41 \pm 0.07$ | 2.30 |
| $Sr^{2+}$ | $0.94 \pm 0.02$ | 0.89 | $1.12 \pm 0.07$ | 0.99 | $1.45 \pm 0.02$ | 1.30 | $1.30 \pm 0.05$ | 2.07 |
| $Ba^{2+}$ | $0.85 \pm 0.01$ | 0.85 | $0.95 \pm 0.02$ | 0.91 | $1.12 \pm 0.01$ | 1.12 | $0.90 \pm 0.04$ | – |

#### S12 Calculation of activity derivatives

Activity derivatives were calculated as described in our previous works.[5,6] Briefly, the activity derivative $a_{cc}$ was calculated using Kirkwood–Buff (KB) theory.[24] KB integrals $G_{ij}$ were obtained from radial distribution functions $g_{ij}(r)$ between species $i$ and $j$:[25]

$$G_{ij} = 4\pi \int_0^\infty \left[g_{ij}(r) - 1\right] r^2\, dr. \tag{S9}$$

Because our simulations were performed in the NPT ensemble with finite box sizes, the integrals were evaluated up to a cutoff radius of 1.99 nm and the radial distribution functions were corrected following the finite-size procedure of Lyubartsev and Laaksonen.[26] The activity derivative is then obtained as[25]

$$a_{cc} = \left(\frac{\partial \ln a_c}{\partial \ln \rho_c}\right)_{p,T} = \frac{1}{1 + \rho_c(G_{cc} - G_{co})}. \tag{S10}$$

with the activity $a_c = \rho_c\gamma_c$, where $\gamma_c$ is the cosolvent molar activity coefficient and $\rho_c$ the number density.

To assess the sensitivity with respect to the finite integration cutoff, the KB integrals and the resulting $a_{cc}$ were evaluated for cutoff radii between 1.50 and 1.99 nm for each trajectory. However, the standard deviation of $a_{cc}$ for this range of cutoffs along each trajectory was found to be significantly smaller than the standard error of the mean over three independent 150 ns trajectories. Therefore, the latter is reported as uncertainties in Tables S18 and S19.

#### S13 Experimental activity derivative

For determining experimental reference values for $a_{cc}$, we first used a quadratic fit of the concentration in molarity (mol/l) as a function of the concentration in molality (mol/kg) based on the data reported in Ref. 27 up to around 2 mol/kg. The equation of this quadratic fit for each salt was used for the conversion from molarity to molality. For the bromide salts, the required data is only reported for the two cations $Na^+$ and $K^+$. So for the other bromide salts, we used the same parameters obtained from the fits of the corresponding chloride salts for the conversion. $a_{cc}$ was then determined via Eq. S10 using mostly the tabulated values of $\gamma_c$ reported in Ref. 28 by employing the central difference scheme for approximating the derivative. The only exceptions are the values for $\gamma_c$ for a concentration of 0.05 mol/kg and that we used $\gamma_c = 0.463$ for a $BaBr_2$ concentration of 0.2 mol/kg, for which we used Ref. 27.

To assess the sensitivity to the numerical differentiation scheme, we also evaluated forward and backward differences in addition to central differences. The maximum deviation between the results from central differences and forward/backward differences is a measure of the numerical differentiation sensitivity and reported as uncertainties for the experimental values. However, this is solely an estimation of the error by numerical differentiation, as no uncertainties for the experimental $a_{cc}$ are reported in Ref. 28.

#### S14 Activity derivatives for different force fields and water models

Table S18: Activity derivatives $a_{cc}$ in OPC water for chloride salts. The current MS/G-LB(OPC) is highlighted.

| | $Li^+$ | $Na^+$ | $K^+$ | $Cs^+$ | $Mg^{2+}$ | $Ca^{2+}$ | $Sr^{2+}$ | $Ba^{2+}$ |
|---|---|---|---|---|---|---|---|---|
| | | | | **$m$ = 0.26 mol/kg** | | | | |
| **MS/G-LB(OPC)** | $0.95 \pm 0.04$ | $0.99 \pm 0.01$ | $0.92 \pm 0.02$ | $0.88 \pm 0.02$ | $0.84 \pm 0.02$ | $0.91 \pm 0.01$ | $0.94 \pm 0.02$ | $0.85 \pm 0.01$ |
| LB/HS | $0.99 \pm 0.01$ | $1.01 \pm 0.01$ | $0.94 \pm 0.02$ | $0.95 \pm 0.04$ | | | | |
| MS | $0.97 \pm 0.02$ | $0.94 \pm 0.03$ | $0.93 \pm 0.02$ | $0.93 \pm 0.01$ | $0.96 \pm 0.02$ | $0.97 \pm 0.03$ | $0.95 \pm 0.03$ | $0.89 \pm 0.03$ |
| LM | $0.96 \pm 0.05$ | $0.94 \pm 0.04$ | $0.88 \pm 0.01$ | $0.89 \pm 0.01$ | $0.88 \pm 0.02$ | $0.87 \pm 0.02$ | $0.89 \pm 0.02$ | $0.88 \pm 0.01$ |
| Exp | $0.96 \pm 0.02$ | $0.92 \pm 0.01$ | $0.89 \pm 0.01$ | $0.86 \pm 0.01$ | $0.93 \pm 0.04$ | $0.90 \pm 0.03$ | $0.89 \pm 0.03$ | $0.85 \pm 0.03$ |
| | | | | **$m$ = 0.52 mol/kg** | | | | |
| **MS/G-LB(OPC)** | $1.02 \pm 0.03$ | $0.97 \pm 0.02$ | $0.94 \pm 0.02$ | $0.87 \pm 0.02$ | $0.96 \pm 0.02$ | $1.10 \pm 0.02$ | $1.12 \pm 0.07$ | $0.95 \pm 0.02$ |
| LB/HS | $1.01 \pm 0.02$ | $0.97 \pm 0.02$ | $0.94 \pm 0.02$ | $0.92 \pm 0.02$ | | | | |
| MS | $1.03 \pm 0.02$ | $1.00 \pm 0.04$ | $0.92 \pm 0.01$ | $0.88 \pm 0.01$ | $1.10 \pm 0.04$ | $1.10 \pm 0.02$ | $1.10 \pm 0.02$ | $0.96 \pm 0.02$ |
| LM | $0.96 \pm 0.01$ | $0.91 \pm 0.02$ | $0.84 \pm 0.02$ | $0.85 \pm 0.02$ | $0.99 \pm 0.01$ | $0.99 \pm 0.02$ | $1.04 \pm 0.01$ | $1.02 \pm 0.03$ |
| Exp | $1.01 \pm 0.03$ | $0.93 \pm 0.01$ | $0.89 \pm 0.01$ | $0.84 \pm 0.01$ | $1.08 \pm 0.04$ | $1.03 \pm 0.04$ | $0.99 \pm 0.03$ | $0.91 \pm 0.01$ |
| | | | | **$m$ = 1.06 mol/kg** | | | | |
| **MS/G-LB(OPC)** | $1.07 \pm 0.01$ | $1.04 \pm 0.03$ | $0.94 \pm 0.01$ | $0.92 \pm 0.01$ | $1.31 \pm 0.04$ | $1.50 \pm 0.07$ | $1.45 \pm 0.02$ | $1.12 \pm 0.01$ |
| LB/HS | $1.15 \pm 0.02$ | $1.09 \pm 0.03$ | $0.97 \pm 0.02$ | $0.98 \pm 0.02$ | | | | |
| MS | $1.17 \pm 0.02$ | $1.05 \pm 0.02$ | $0.95 \pm 0.04$ | $0.90 \pm 0.05$ | $1.54 \pm 0.04$ | $1.56 \pm 0.02$ | $1.57 \pm 0.04$ | $1.06 \pm 0.02$ |
| LM | $0.99 \pm 0.02$ | $0.94 \pm 0.03$ | $0.84 \pm 0.01$ | $0.84 \pm 0.02$ | $1.34 \pm 0.02$ | $1.34 \pm 0.03$ | $1.46 \pm 0.05$ | $1.39 \pm 0.02$ |
| Exp | $1.15 \pm 0.03$ | $0.97 \pm 0.01$ | $0.90 \pm 0.01$ | $0.84 \pm 0.01$ | $1.53 \pm 0.08$ | $1.38 \pm 0.07$ | $1.30 \pm 0.05$ | $1.12 \pm 0.03$ |
| | | | | **$m$ = 2.12 mol/kg** | | | | |
| **MS/G-LB(OPC)** | $1.30 \pm 0.03$ | $1.12 \pm 0.01$ | $0.97 \pm 0.02$ | $0.96 \pm 0.03$ | $2.08 \pm 0.11$ | $0.41 \pm 0.07$ | $1.30 \pm 0.05$ | $0.90 \pm 0.04$ |
| LB/HS | $1.47 \pm 0.04$ | $1.33 \pm 0.03$ | $1.08 \pm 0.03$ | $1.15 \pm 0.05$ | | | | |
| MS | $1.36 \pm 0.01$ | $1.11 \pm 0.02$ | $1.01 \pm 0.02$ | $0.97 \pm 0.02$ | $3.01 \pm 0.03$ | $0.38 \pm 0.06$ | $1.34 \pm 0.09$ | $0.69 \pm 0.06$ |
| LM | $1.19 \pm 0.04$ | $1.01 \pm 0.04$ | $0.71 \pm 0.03$ | $0.86 \pm 0.02$ | $1.93 \pm 0.07$ | $0.16 \pm 0.02$ | $1.16 \pm 0.09$ | $2.37 \pm 0.12$ |
| Exp | $1.47 \pm 0.08$ | $1.12 \pm 0.04$ | $0.96 \pm 0.02$ | $0.89 \pm 0.01$ | $2.67 \pm 0.29$ | $2.30 \pm 0.23$ | $2.07 \pm 0.21$ | – |

Table S19: Activity derivatives $a_{cc}$ in OPC water for bromide salts. The current MS/G-LB(OPC) is highlighted.

| | $Li^+$ | $Na^+$ | $K^+$ | $Cs^+$ | $Mg^{2+}$ | $Ca^{2+}$ | $Sr^{2+}$ | $Ba^{2+}$ |
|---|---|---|---|---|---|---|---|---|
| | | | | | **$m$ = 0.26 mol/kg** | | | |
| **MS/G-LB(OPC)** | $1.00 \pm 0.01$ | $0.99 \pm 0.03$ | $0.91 \pm 0.02$ | $0.87 \pm 0.02$ | $0.87 \pm 0.02$ | $0.97 \pm 0.03$ | $0.98 \pm 0.02$ | $0.97 \pm 0.02$ |
| LB/HS | $0.97 \pm 0.02$ | $0.98 \pm 0.02$ | $0.97 \pm 0.03$ | $0.95 \pm 0.04$ | | | | |
| LM | $0.97 \pm 0.02$ | $0.95 \pm 0.04$ | $0.90 \pm 0.01$ | $0.89 \pm 0.02$ | $0.95 \pm 0.01$ | $0.95 \pm 0.01$ | $0.96 \pm 0.02$ | $0.95 \pm 0.01$ |
| Exp | $0.96 \pm 0.02$ | $0.92 \pm 0.01$ | $0.90 \pm 0.01$ | $0.86 \pm 0.02$ | $0.98 \pm 0.05$ | $0.94 \pm 0.05$ | $0.92 \pm 0.04$ | $0.90 \pm 0.06$ |
| | | | | | **$m$ = 0.52 mol/kg** | | | |
| **MS/G-LB(OPC)** | $1.03 \pm 0.01$ | $1.01 \pm 0.03$ | $0.91 \pm 0.03$ | $0.90 \pm 0.02$ | $0.99 \pm 0.04$ | $1.17 \pm 0.02$ | $1.16 \pm 0.02$ | $1.15 \pm 0.03$ |
| LB/HS | $1.08 \pm 0.02$ | $1.03 \pm 0.03$ | $0.95 \pm 0.02$ | $0.92 \pm 0.02$ | | | | |
| LM | $1.04 \pm 0.03$ | $0.97 \pm 0.03$ | $0.87 \pm 0.01$ | $0.84 \pm 0.01$ | $1.09 \pm 0.02$ | $1.17 \pm 0.02$ | $1.15 \pm 0.03$ | $1.13 \pm 0.01$ |
| Exp | $1.02 \pm 0.02$ | $0.96 \pm 0.01$ | $0.90 \pm 0.01$ | $0.83 \pm 0.01$ | $1.20 \pm 0.07$ | $1.11 \pm 0.05$ | $1.04 \pm 0.03$ | $1.01 \pm 0.04$ |
| | | | | | **$m$ = 1.06 mol/kg** | | | |
| **MS/G-LB(OPC)** | $1.26 \pm 0.02$ | $1.10 \pm 0.05$ | $0.99 \pm 0.01$ | $0.92 \pm 0.01$ | $1.36 \pm 0.05$ | $1.75 \pm 0.06$ | $1.85 \pm 0.03$ | $1.64 \pm 0.04$ |
| LB/HS | $1.27 \pm 0.03$ | $1.15 \pm 0.02$ | $1.03 \pm 0.02$ | $0.98 \pm 0.02$ | | | | |
| LM | $1.19 \pm 0.04$ | $1.10 \pm 0.04$ | $0.82 \pm 0.02$ | $0.81 \pm 0.02$ | $1.55 \pm 0.07$ | $1.58 \pm 0.01$ | $1.64 \pm 0.03$ | $1.71 \pm 0.03$ |
| Exp | $1.21 \pm 0.04$ | $1.03 \pm 0.03$ | $0.92 \pm 0.01$ | $0.84 \pm 0.01$ | $1.76 \pm 0.12$ | $1.56 \pm 0.08$ | $1.44 \pm 0.08$ | $1.28 \pm 0.06$ |
| | | | | | **$m$ = 2.12 mol/kg** | | | |
| **MS/G-LB(OPC)** | $1.58 \pm 0.04$ | $1.35 \pm 0.04$ | $0.99 \pm 0.03$ | $0.92 \pm 0.01$ | $2.28 \pm 0.11$ | $0.42 \pm 0.04$ | $1.80 \pm 0.07$ | $2.12 \pm 0.16$ |
| LB/HS | $1.53 \pm 0.04$ | $1.43 \pm 0.03$ | $1.10 \pm 0.04$ | $1.15 \pm 0.05$ | | | | |
| LM | $1.60 \pm 0.06$ | $1.21 \pm 0.03$ | $0.74 \pm 0.03$ | $0.78 \pm 0.01$ | $2.73 \pm 0.03$ | $0.45 \pm 0.04$ | $1.42 \pm 0.12$ | $2.70 \pm 0.11$ |
| Exp | $1.59 \pm 0.09$ | $1.22 \pm 0.04$ | $0.99 \pm 0.03$ | $0.87 \pm 0.02$ | $3.09 \pm 0.33$ | $2.73 \pm 0.28$ | – | – |

#### S15 Activity derivative for bromide salts for MS/G-LB(OPC)

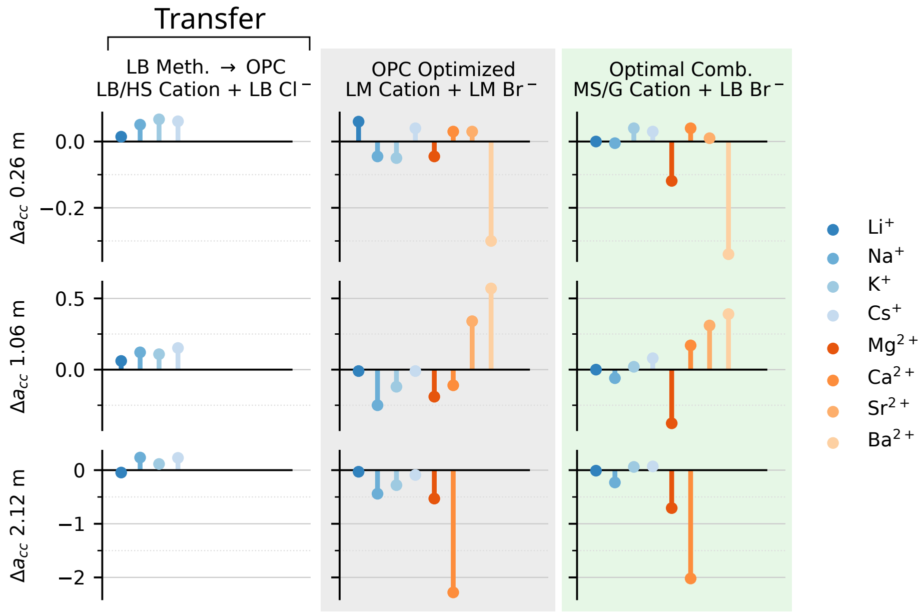


Figure S4: **Difference between simulated and experimental activity derivatives:** $\Delta a_{cc}$ at 0.26 m (top), at 1.06 m (middle) and 2.12 m (bottom). Experimental values were obtained from the data reported in Ref. 28.

### S16 Radial distribution functions for MS/G-LB(OPC)

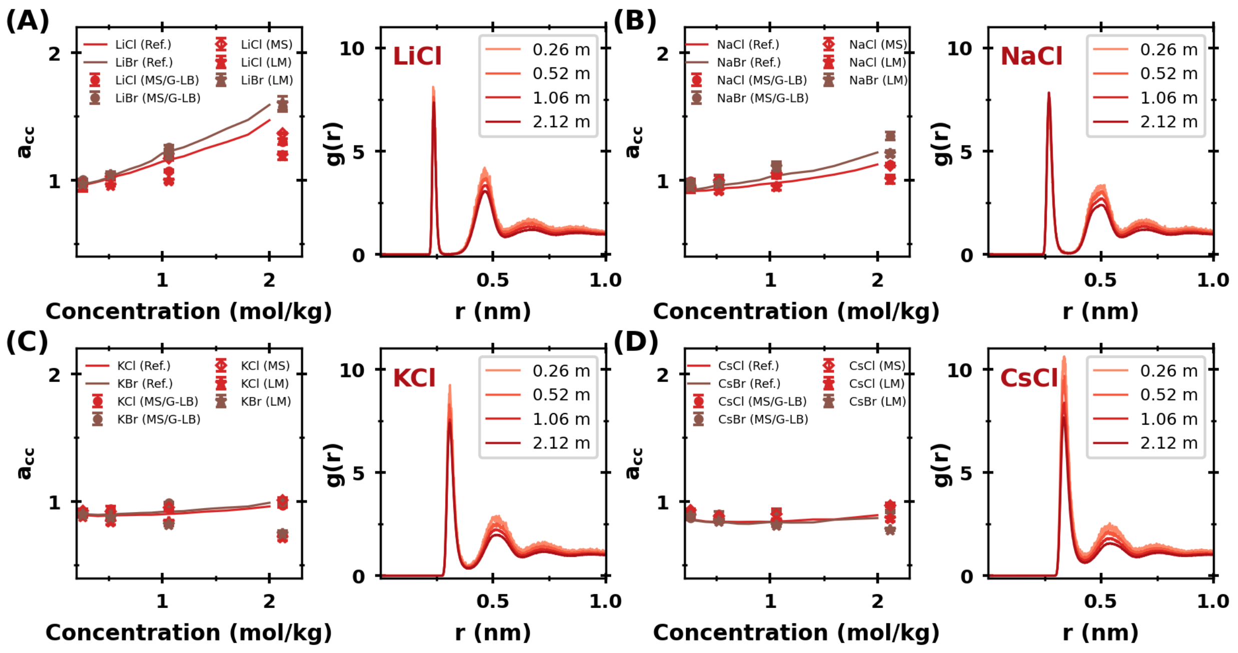


Figure S5: **Activity derivatives ($a_{cc}$) and radial distribution functions (RDF) for MS/G-LB(OPC), MS(OPC), LM(OPC):** (A) LiCl/Br, (B) NaCl/Br, (C) KCl/Br and (D) CsCl/Br. $a_{cc}$ is shown as function of concentration in mol/kg and for reference values determined from experimental data.[27,28] RDFs between cation and chloride for four different concentrations are shown for the MS/G-LB(OPC) parameters.

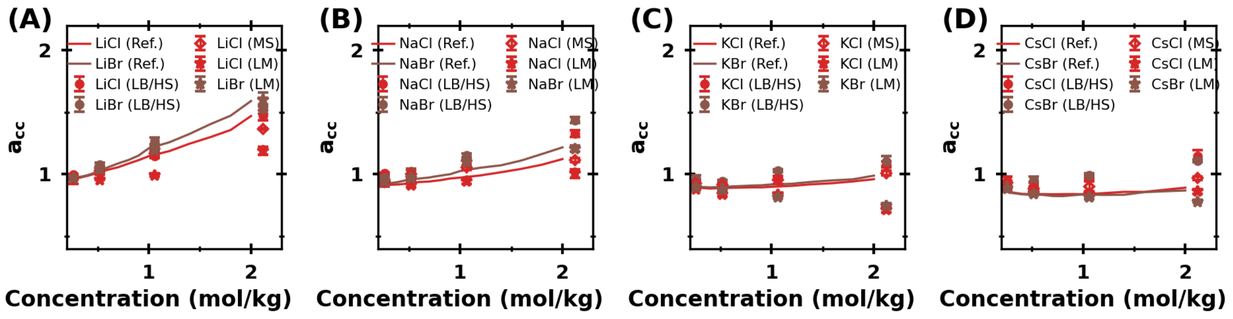


Figure S6: **Activity derivatives ($a_{cc}$) and radial distribution functions (RDF) for LB/HS(OPC), MS(OPC), LM(OPC).**

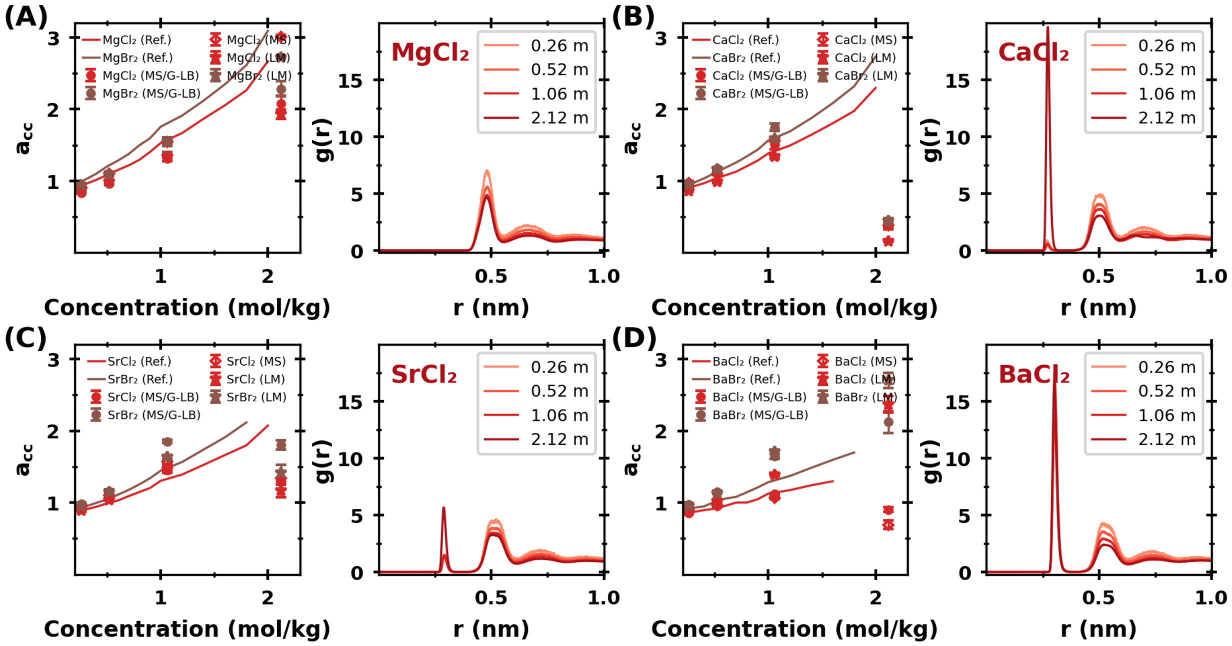


Figure S7: **Activity derivatives ($a_{cc}$) and radial distribution functions (RDF) for MS/G-LB(OPC), MS(OPC), LM(OPC):** (A) $MgCl_2/Br_2$, (B) $CaCl_2/Br_2$, (C) $SrCl_2/Br_2$ and (D) $BaCl_2/Br_2$. $a_{cc}$ is shown as function of concentration in mol/kg and for reference values determined from experimental data.[27,28] RDFs between cation and chloride for four different concentrations are shown for the MS/G-LB(OPC) parameters.

#### S17 Calculation of self-diffusion coefficients

The diffusion coefficients were calculated as in our previous work.[5,6] Briefly, the diffusion coefficients were obtained from additional 50 ns NVT simulations. Diffusion coefficients were determined from the slope of the mean-square displacement using the Einstein relation. The diffusion coefficients obtained under periodic boundary conditions were corrected for finite-size effects following Yeh and Hummer:[29]

$$D_0 = D_{\mathrm{pbc}}(L) + \frac{k_B T \xi_{\mathrm{ew}} \alpha}{6\pi \eta_{\mathrm{Model}} L}. \tag{S11}$$

where $D_0$ is the diffusion coefficient in the infinite-system limit, $D_{\mathrm{pbc}}(L)$ the value obtained from the simulation box of length $L$, $k_B$ the Boltzmann constant, $T$ the temperature, $\xi_{\mathrm{ew}} = 2.837297$ the Ewald self-term for a cubic lattice, $\alpha = 1$, and $\eta_{\mathrm{Model}}$ the viscosity of the corresponding water model. To account for differences between the viscosities of the employed water models and experimental water, the diffusion coefficients were rescaled as

$$D = \frac{\eta_{\mathrm{Model}}}{\eta_{\mathrm{Exp}}} D_0. \tag{S12}$$

The viscosities used were $\eta_{\mathrm{TIP3P}} = 3.21 \times 10^{-4}\,\mathrm{kg\,m^{-1}\,s^{-1}}$ and $\eta_{\mathrm{SPC/E}} = 7.29 \times 10^{-4}\,\mathrm{kg\,m^{-1}\,s^{-1}}$,[30] and $\eta_{\mathrm{OPC}} = 8.0 \times 10^{-4}\,\mathrm{kg\,m^{-1}\,s^{-1}}$.[31] The experimental viscosity of water was taken as $\eta_{\mathrm{Exp}} = 8.91 \times 10^{-4}\,\mathrm{kg\,m^{-1}\,s^{-1}}$.[30]

Reported errors correspond to the standard error of the mean from three independent simulations. Experimental reference values were taken from Ref. 15.

#### S18 Self-diffusion coefficient for different force fields and water models

Table S20: Self-diffusion coefficient $D$ at infinite dilution for monovalent and divalent cations as well as $Cl^-$ and $Br^-$. The current MS/G-LB(OPC) is highlighted. Experimental values were taken from Ref. 15.

| | $Li^+$ | $Na^+$ | $K^+$ | $Cs^+$ | $Mg^{2+}$ | $Ca^{2+}$ | $Sr^{2+}$ | $Ba^{2+}$ |
|---|---|---|---|---|---|---|---|---|
| MS (TIP3P) | 1.16 ± 0.06 | 0.86 ± 0.07 | 1.38 ± 0.07 | 1.37 ± 0.10 | – | 0.80 ± 0.03 | 0.75 ± 0.03 | 0.71 ± 0.03 |
| **MS/G (OPC)** | 1.20 ± 0.01 | 0.93 ± 0.02 | 1.50 ± 0.09 | 1.68 ± 0.08 | – | 0.83 ± 0.04 | 0.76 ± 0.03 | 0.74 ± 0.04 |
| FN/MN (SPC/E) | – | 1.12 ± 0.03 | 1.57 ± 0.04 | 1.78 ± 0.11 | – | 0.82 ± 0.03 | 0.73 ± 0.04 | 0.74 ± 0.03 |
| FN/MN (OPC) | – | 1.08 ± 0.04 | 1.46 ± 0.06 | 1.62 ± 0.04 | – | 0.80 ± 0.04 | 0.71 ± 0.03 | 0.70 ± 0.04 |
| LB/HS (SPC/E) | 1.15 ± 0.04 | 1.14 ± 0.04 | 1.62 ± 0.10 | 1.70 ± 0.17 | – | – | – | – |
| LB/HS (OPC) | 1.26 ± 0.06 | 0.93 ± 0.03 | 1.41 ± 0.05 | 1.55 ± 0.05 | – | – | – | – |
| LM | 1.06 ± 0.08 | 0.99 ± 0.03 | 1.44 ± 0.03 | 1.84 ± 0.08 | – | 0.78 ± 0.05 | 0.71 ± 0.02 | 0.72 ± 0.03 |
| Exp | 1.029 | 1.334 | 1.957 | 2.056 | – | 0.792 | 0.791 | 0.847 |

| | $Cl^-$ | $Br^-$ |
|---|---|---|
| MS (TIP3P) | 1.45 ± 0.09 | – |
| MS (OPC) | 1.48 ± 0.09 | – |
| FN/MN (SPC/E) | 1.60 ± 0.11 | 1.55 ± 0.03 |
| FN/MN (OPC) | 1.65 ± 0.11 | 1.44 ± 0.07 |
| LB/HS (SPC/E) | 1.37 ± 0.03 | 1.56 ± 0.02 |
| **LB/HS (OPC)** | 1.45 ± 0.09 | 1.55 ± 0.02 |
| LM | 1.78 ± 0.11 | 1.80 ± 0.03 |
| Exp | 2.032 | 2.08 |
